# Memory Consolidation with Orthogonal Gradients for avoiding Catastrophic Forgetting

**DOI:** 10.1101/2022.02.25.481890

**Authors:** Tamizharasan Kanagamani, Rupak Krishnamurthy, V. Srinivasa Chakravarthy, Balaraman Ravindran, Ramshekhar N. Menon

## Abstract

The memory consolidation process enables the accumulation of recent and remote memories in the long-term memory store. In general, the deep network models of memory suffer from forgetting old information while learning new information, called catastrophic forgetting/interference. The human brain overcomes this problem quite effectively, a problem that continues to challenge current deep neural network models.

We propose a regularization-based model to solve the problem of catastrophic forgetting. According to the proposed training mechanism, the network parameters are constrained to vary in a direction orthogonal to the average of the error gradients corresponding to the previous tasks. We also ensure that the constraint used in parameter updating satisfies the locality principle.

The proposed model’s performance is compared with Elastic Weight Consolidation on standard datasets such as permuted MNIST and split MNIST on classification tasks using fully connected networks, and Convolution-based networks. The model performance is also compared to an autoencoder on split MNIST dataset, and to complex core50 dataset on two types of classification tasks with EWC.

The proposed model gives a new view on plasticity at the neuronal level. In the proposed model, the parameter updating is controlled by the neuronal level plasticity rather than synapse level plasticity as in other standard models. The biological plausibility of the proposed model is discussed by linking the extra parameters to synaptic tagging, which represents the state of the synapse involved in Long Term Potentiation (LTP).

## Introduction

The human brain continuously acquires, processes, and uses information from the world throughout its lifetime. This lifelong learning is crucial for human survival and effective manipulation in the world since the organism needs to select and remember details of the world that enhance or compromise its existence. The process of formation of memories, particularly over the longer term, known as memory consolidation, predominantly occurs during sleep [1], [2]. Memory consolidation is a time-dependent process in which the recently acquired important information is transferred from the short-term memory and integrated and stabilized in the long-term memory [3]. The importance can be attributed to various factors such as novelty, familiarity, and saliency [4]–[6]. In the consolidation process, the newly acquired memory is enhanced and integrated with the previously existing memories in the long-term store [7].

Squire et al. (1994) proposed the theory of standard memory consolidation, in which the hippocampus serves as a buffer, a “scratch pad” (temporary store). During consolidation, the information from the hippocampus is transferred to the neocortex (permanent store) [8]. On the other hand, multiple trace theory proposes that episodic memory is initially dependent on the hippocampus, and after consolidation, it depends on both the hippocampus and the neocortex. In contrast, the semantic memory depends only on the neocortex[9]–[11].

In episodic memory formation, both the hippocampus and the prefrontal cortex play a major role in the consolidation process [12]. This consolidation is initiated by the process of replaying the sequences of acquired memory in the hippocampus during the slow-wave stages of sleep [13], [14]. Miller and Cohen explained that episodic memory processing involves the interaction between the hippocampus and the medial prefrontal cortex (mPFC) [15], and this interaction is supported by the bidirectional projections [16]. The encoding and the maintenance of the memory involve the flow of information from the hippocampus to the mPFC [16]–[18], whereas memory retrieval involves the flow of information from the mPFC to the hippocampus [16]. Cooper et al. described the network between the sensory cortex and the hippocampus as the short-term memory, and the network between the sensory-cortex and the prefrontal cortex as the long-term memory [19], [20]. Short-term memory has a lesser capacity and can retain only less information; on the other hand, the long-term memory has a higher capacity and can retain older information alongside the new information [21]. This retention is applicable even when the old information is not visited again.

Several biologically inspired deep neural network models have been proposed to explain this memory retention behavior in the long-term memory. But none of these models can match the capability of the human brain. These models fail significantly in retaining the old information during training with new information. This behavior is referred to as catastrophic forgetting or catastrophic interference [22].

A variety of approaches have been proposed to solve the problem of catastrophic forgetting. Some approaches replay the complete or portion of the old data while training the new data. Robins et al. proposed a model that uses an episodic memory system that stores the old data [23]. Since old data from the episodic memory is trained along with the new data, this method needs a large memory space for storing old data. Shin et al. proposed a deep generative model where a generator creates samples from the previous tasks, and the generated samples are trained together with the new data [24]. Van de Ven et al proposed a replay-based model, where the internal representations are replayed by context specific generator model [25] .

Some approaches use the expansion of networks at multiple layers based on certain criteria. Rusu et al. proposed the Progressive Network, which creates a separate neural network (column) for each task [26], wherein transfer is allowed through lateral connections to the features of already existing neural networks (columns). In this approach, when training one task, network parameters corresponding to the previous task are fixed.

Self-net trains a separate network for each task and stores the trained parameters for each task using an autoencoder in a compressed format. By using the latent variables, the parameters for a particular task can be retrieved from the autoencoder and reused [27].

Yoon et al. proposed dynamically expandable networks [28]. This type of network selects the neurons appropriate to the new task and allows only the parameters corresponding to those neurons to learn. If this selective training fails to learn the new task, then the number of neurons is increased in a top-down manner.

Masse et al. proposed a context-dependent gating method, where the network uses context signals along with the inputs. In this method, the network parameters are split at each layer for each context. In another variant of this approach, 80% of parameters are gated, and only 20% of the parameters are updated for a particular task [29].

Some methods use controlled learning of network parameters. Li and Hoem proposed learning without forgetting method, where the network uses two sets of parameters: shared parameters and task-specific parameters [30]. The shared parameters include all the parameters except the task-specific output layer parameters. This network only uses the new task data for preserving the network performance on earlier tasks. Before training the network on a new task, the outputs for the new task data on each old task networks are estimated. During training on the new task, both the new task network loss and the old task network loss on the output are minimized together.

Kirkpatrick et al. proposed Elastic Weight Consolidation (EWC), where the network parameter change is controlled with the help of the diagonals of the Fisher information matrix corresponding to the previous tasks [31]. This approach keeps a copy of the diagonals of the Fisher information matrix and the latest learned parameters. Zenke et al. proposed the Synaptic Intelligence model, where the importance of each parameter for the previous tasks is estimated based on the change in parameter values [32]. Based on this value, the parameters with high importance refrain from learning, and the rest of the parameters are allowed to update. In Synaptic Intelligence [32], the importance is estimated online fashion, whereas in EWC, the diagonals of the Fisher information matrix are estimated after training each task. Farajtabar et al. (2019) proposed the orthogonal gradient descent method, where the gradients of the new task are projected to a subspace in which the gradients do not affect the output of the old tasks [33]. This method maintains a fixed number of gradients for each task and uses Gram-Schmidt process for finding the orthogonal direction.

Schwarz et al. proposed the progress and compress model, where two neural networks (active column and knowledge base) are used and trained in two phases [34]. During the progress phase, only the active column is trained with the new task. The connections between layers of the active column and knowledge base are established to enhance the reuse of encoded features as it allows the transfer of old tasks. During the compression phase, the active column is transferred to the knowledge base using the distillation process as proposed by Hinton et al. [35]. Maltoni et al. introduced the AR1 model, which proposes the use of regularization methods along with network expansion methods for single-incremental-tasks [36].

In this work, we propose a model to solve the problem of catastrophic forgetting by controlling the direction in which the parameters are updated. This is realized by adding an extra regularization-based loss function along with a conventional loss function. This extra loss function restricts the parameter updation in some directions which is important for the earlier learned tasks. While the other models use the constraints at the connection/synapse level or at the network level, the proposed model provides constraints at the neuronal level. EWC and SI are state-of-the-art approaches showing higher performance among the regularization-based methods. Therefore, the performance of the proposed model is compared with the elastic weight consolidation (EWC) model alone. For evaluating the performance, MNIST and Core50 datasets are used. MNIST is a dataset containing handwritten numbers (0 - 9). Core50 is a dataset developed for solving continual learning problems with real-world objects. A detailed description of how the datasets are used is explained in the dataset section. We verify if the proposed model can retain old information upon training with the new information in explaining the memory consolidation behavior. The performance has been evaluated on MultiLayer Perceptron (MLP) based classification tasks on the permuted MNIST dataset and split MNIST dataset. The performance has also been evaluated on Autoencoder and convolution-based classification tasks using the split MNIST dataset. The performance on complex datasets is evaluated using the CORe50 dataset [37] on two types of tasks, namely New Instances (NI) and New Classes (NC).

## Methods and Results

To solve the problem of catastrophic forgetting in a continual learning setting, we proposed a method in which the parameters are updated by constraining the direction of update. Instead of giving importance to individual parameters just as in earlier methods like EWC and SI, the proposed model gives more preference to a particular direction for the update of the input parameters corresponding to a specific neuron. Here the parameters are constrained to move orthogonally to the previous direction. The need for this controlled update of parameters and the implementation method is described below.

To justify this special constraint and to define the way to update the parameters for a task, it is convenient to consider neural network training from a linear perspective.

Consider the below linear regression equation (a single neuron with linear activation function).

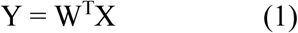

Let X ∈ R^m^ be the input, W∈ R^m^ be the parameters, and Y∈ R^1^ be the output.

### Theorem 1

Let *Y*_1_ = *W*^*T*^*X*_1_ and *W*^i^ = *W* + Δ*W*. Here Δ*W*∈ R^m^ is the change in W. There exists W′ which satisfies *Y*_1_ = *W*′^*T*^*X*_1_ if and only if Δ*W* ⊥ *X*_1_.

. In the above linear regression problem, consider the loss function 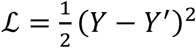, where X is the input vector, W is the weight vector (parameters), Y is the expected output and Y’= W^*T*^X is the estimated output.

### Proposition 1

If 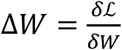 is the gradient vector, then both the X and Δ*W* vectors represent the same unsigned direction.

### Proof

Consider the first-order derivative of the loss with respect to W:

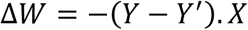

It can be observed that X and Δ*W* are different in terms of magnitude. But they denote the same unsigned direction. This is valid for any activation function.

For example, if sigmoid activation function is used then Y = *σ* (X^T^W) and the first-order derivative will be Δ*W* = (*Y* − *Y*^*i*^). *Y*^*i*^. (1 − *Y*^i^). *X*.

By combining the above two observations, we can conclude that, if the change in the parameters Δ*W* (while training the second data), is orthogonal to the gradient corresponding to the first data, Δ*W*_1_, then the representations generated from the first data will remain intact upon updating. This is reminiscent of null space control when combining controllers in classical control systems. If the second controller is in the null space of the first one, then the first control’s stability/performance is not compromised.

This idea can be extended to any deep neural network by restricting the change in parameters to the orthogonal direction to the input vector. In deep neural networks, the inputs of each neuron can be related to the gradients of its input parameters.

To summarize, when the network is trained on the second dataset, if the parameters change in the orthogonal direction to the gradients of the first data at the neuronal level, then the representations of the first data used to train the network can be retained. This constraint can be accomplished by adding the following term to the loss function.

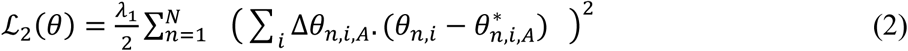

where *ℒ*_2_ is the extra loss term used along with standard loss; Δ*θ*_*n,i,A*_ is the gradient for the first data set A; *θ*_*n,i*_ represents the current network parameter; 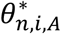 represents the trained parameters for the data A; n represents individual neuron and ‘i’ represents each input parameter to that particular neuron.

When the network is trained using this approach, the copy of the trained parameters (*θ*^∗^) and the gradients (Δ*θ*) needs to be preserved. Thus, the network maintains three times the number of actual parameters. As the network is trained in batches, after training the first batch, the average of the normalized gradients is estimated and used for further training. Here normalization is employed to give equal importance for all the data points in the batch.

To evaluate the proposed model, the performance on alleviating catastrophic forgetting is tested using deep neural networks. In particular, a network of four fully connected layers is trained on a sequence of classification tasks. Once the network is trained on a particular task, the network is not allowed for further training on any data from the same task. The model is evaluated on the permuted MNIST dataset and split MNIST dataset. The performance is also evaluated on complex networks such as CNN and Autoencoder using split MNIST datasets. The performance is also evaluated with the complex dataset CORe50. Although some methods perform better than EWC in continual learning settings, the realization of biological plausibility is not observed in these models. Some approaches keep copies of task-related meta-data, such as gradients for each task. These meta-data storage increase as the number of tasks increases [33]. The replay-based approaches keep the subset of earlier trained data [38]–[40] or use a generator network for sampling from earlier tasks [24]. Among the different methods, the proposed model uses a similar regularization-based mechanism as in EWC and SI. They also explain biological plausibility in terms of synaptic plasticity. So the performance of the proposed model is compared with that of EWC [31].

## Dataset

### Permuted MNIST

For the permuted MNIST task, 1000 images are chosen from the MNIST dataset as task 1 (T1). For each task, one fixed random permutation is chosen and applied to the input pixels (784) of all the images. In this manner, three tasks are created, each having 1000 images.

### Split MNIST

For the split MNIST, the MNIST images are divided into three tasks, each having only three classes. In each task, the network has to predict one among the three classes. Each task has the images of classes [0,1,2], [3,4,5] and [6,7,8] respectively. Here also, for each task, 1000 images are chosen. The data distribution for the split MNIST dataset is shown in Table 1.

**Table 1.** Split MNIST Task distribution

| Task No | Images from MNIST Dataset |
| --- | --- |
| Task 1 | 0,1,2 |
| Task 2 | 3,4,5 |
| Task 3 | 6,7,8 |

### Network Architecture

The network comprises 4 fully connected layers. It receives the input of size 784 and the data is processed by fully connected layers with 256, 128, 64, 10 neurons respectively. For the split MNIST tasks, the last layer of the network uses 3 neurons. For both the tasks, the first two layers use the rectified Linear Unit (ReLU) activation function. The third layer uses sigmoid function and the final layer uses the Softmax activation function.

### Training

For the first task (A), the network is trained to minimize the conventional cross-entropy loss along with L_2_ regularization loss (Eqn. 4).

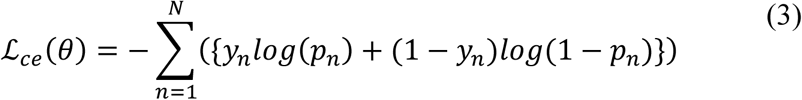

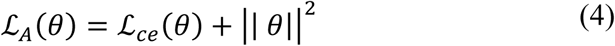

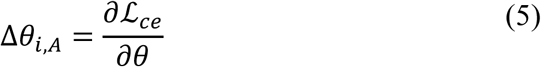

where *ℒ*_*A*_ is the loss function for the first task (A); *ℒ*_*ce*_ is the cross-entropy loss; y_n_ is the true probability of n^th^ class, and p_n_ is the estimated probability for the n^th^ class; N denotes the number of classes; *θ* represents network parameters. For permuted MNIST dataset N = 10 and for split MNIST dataset N=3.

After training the network with the first task (A), a copy of the trained parameters 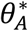 and the corresponding normalized mean gradients Δ*θ*_*i,A*_ are retained and used for further training. These gradients are estimated for task A. These mean gradients are estimated as the average of normalized gradients. This normalization of gradient vector is implemented at the level of neurons.

For the second task, the network is trained with the loss function defined in equation 6. The third task is trained with the loss function given in equation 7.

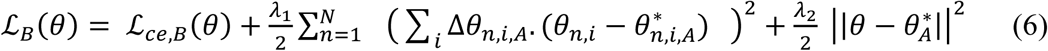

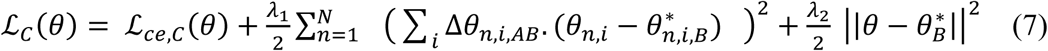

where, *ℒ*_*B*_, *ℒ*_*C*_ are the loss functions used for the tasks B, and C in the proposed model; *ℒ*_*ce,B*_(*θ*) is the cross-entropy loss for task B; *ℒ*_ce,C_(*θ*) is the cross-entropy loss for task C; *θ* represents the network parameters; *θ*^∗^ is the latest trained parameter for the previous task. Δ*θ*_*i,A*_ is the gradient of cross-entropy to the trained parameters 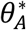. Δ*θ*_*i,AB*_ is the average of the gradients corresponding to the tasks A and B. Thus, when the network is trained on multiple tasks, Δθ is estimated as the average of all the gradients from all the trained tasks. *λ*_1_, *λ*_2_ are the trade-off parameters. Here the second term in the loss function restricts the movement of parameters to the orthogonal direction of Δ*θ*_*i,A*_ and the third term is used to keep the parameters as close as possible to the earlier trained parameters, thus acts as a regularization term. In all the cases, the network is trained using Adam optimizer. From the second task onwards, Δ*θ* is estimated as the average of the normalized gradients for all the previously trained tasks.

The performance of the proposed model on retaining the old information is compared with EWC. In the EWC, the network is trained to minimize the loss function given in Eqn. 8.

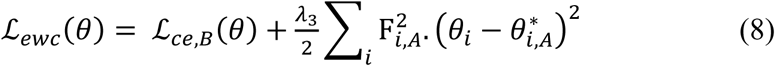

where *ℒ*_ewc_ is the loss function used by the EWC. 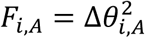 represent the diagonal elements of Fisher information matrix estimated from the first-order derivatives of the loss function with respect to the network parameters for task A, *λ*_3_ is the trade-off parameter.

## Results

The performance of the proposed method is compared with EWC on both the permuted MNIST and Split MNIST tasks using fully connected deep networks. Here both the methods use the same network architecture. In order to evaluate the proposed method on complex networks, a split MNIST dataset alone is used on CNN-based classifier and Autoencoder. For evaluating a complex dataset, the CORe50 dataset is used.

### Permuted MNIST

From the MNIST dataset, three permuted tasks, each with 1000 data samples, are created as described earlier. The first task, T1, contains MNIST images. The remaining tasks, T2 and T3, contain two different permuted versions of T1. Here the tasks are trained in the order T1, T2, and T3. Each task is trained for a fixed number of epochs. Once the network is trained for a particular task, it will not receive any data from that task again.

The network is trained on the first task T1 to 100% accuracy. When the network is trained on tasks T2 and T3 in order, the accuracy on each task is compared between the proposed model and EWC.

Figure 2 shows the accuracy on the tasks T1 (red), T2 (green), and T3 (blue) while training using EWC. The X-axis shows the training stages. At each stage, the network is trained on only one task. At the end of the training, the tasks T1, T2, and T3 show an accuracy of 98.7%, 99.5%, and 98.9%, respectively. Thus, EWC attains an average accuracy of 99.03%.

**Figure 1.**
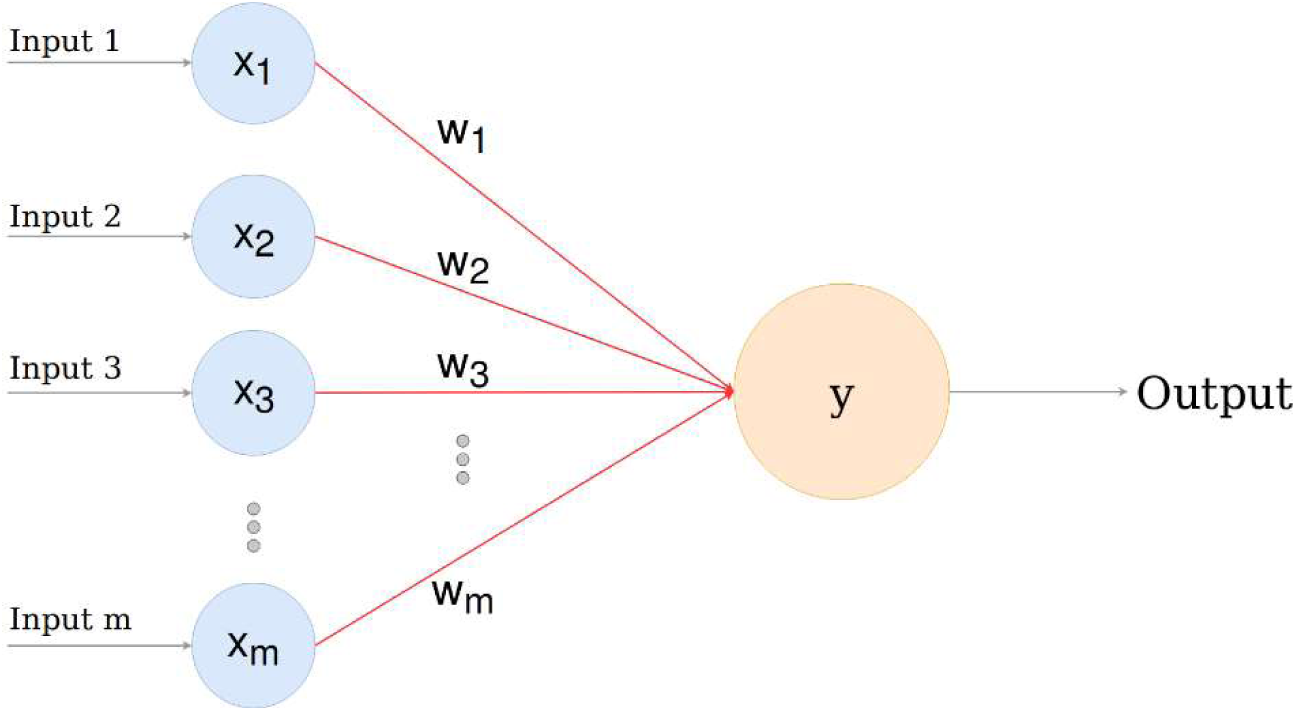
Linear Regression

**Figure 2.**
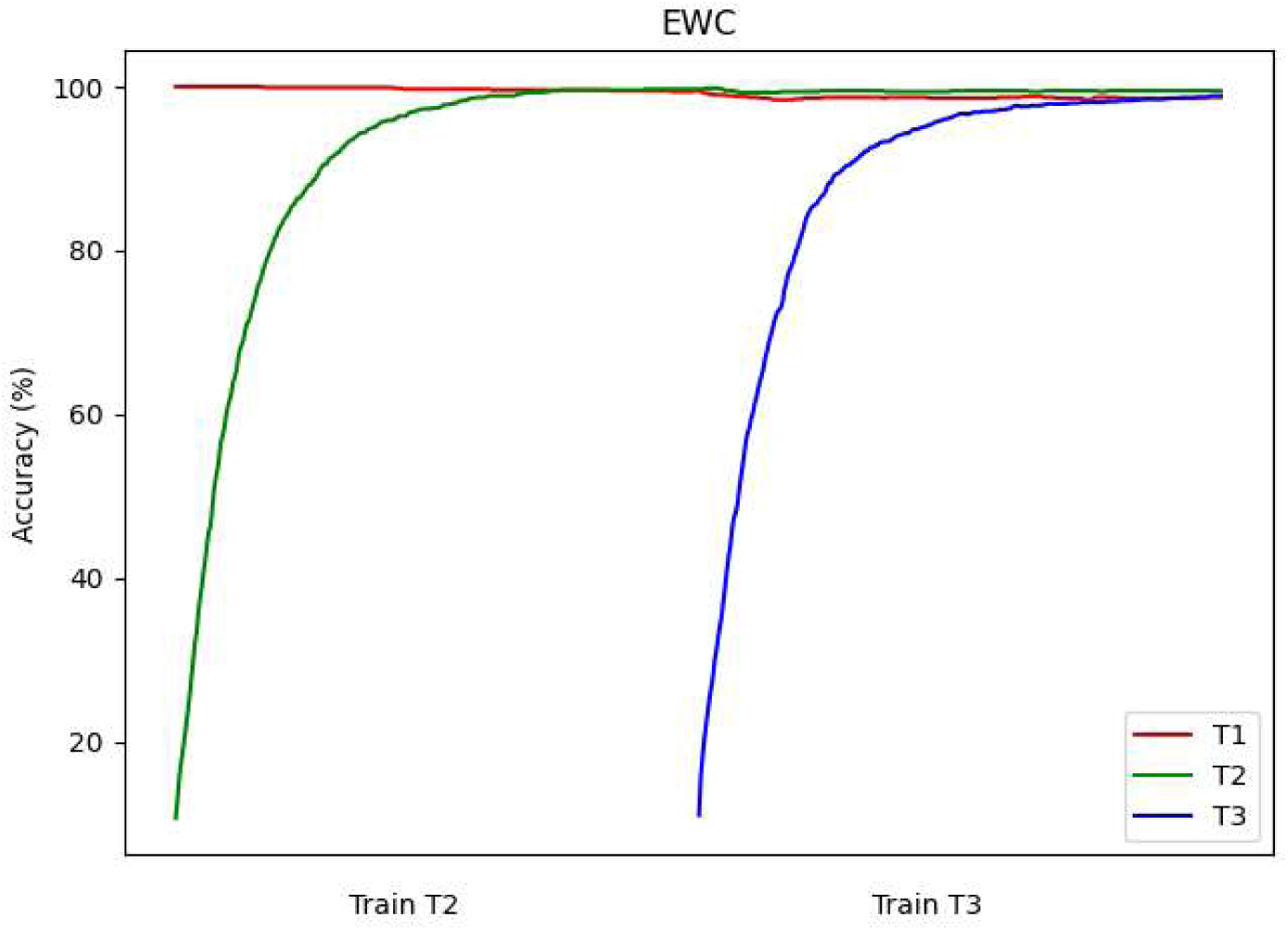
Task accuracies on permuted MNIST Dataset using EWC

Figure 3 shows the accuracy on the task T1 (red), T2 (green), and T3 (blue) while training using the proposed model. After training all the tasks, T1, T2, and T3 get the accuracy of 99.3%, 98.6%, and 100%, respectively. The proposed model gets an average accuracy of 99.3%.

**Figure 3.**
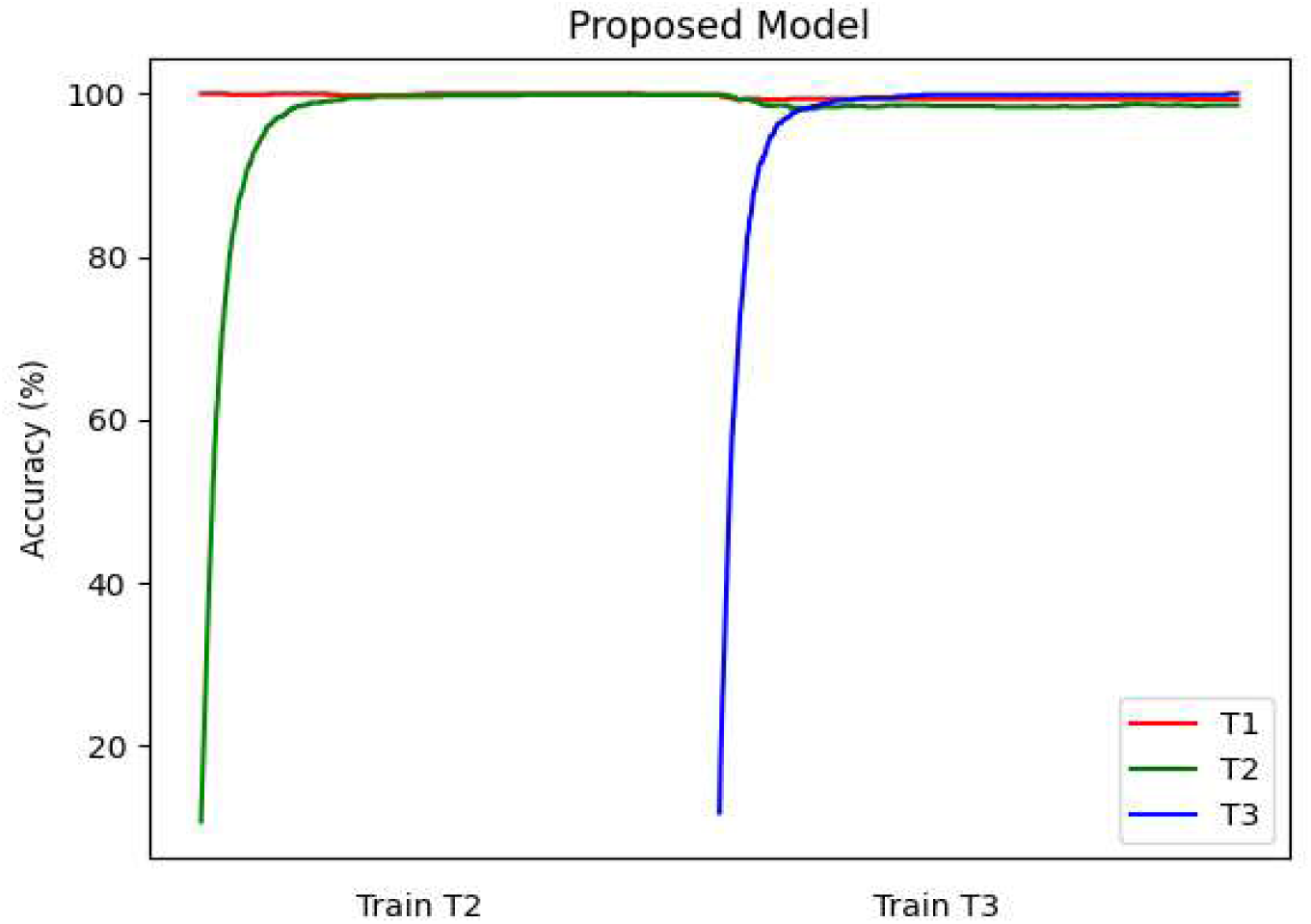
Task accuracies on permuted MNIST Dataset using the proposed model

### Split MNIST

Here the MNIST dataset is split into three tasks (T1, T2, and T3), each having 1000 images and, each image corresponds to one among three classes as described before. Here also, the first task, T1, is trained to 100% accuracy, and the level of retaining the accuracy for all the tasks is plotted for the EWC and the proposed model while training on the tasks T2 and T3.

Figure 4 shows the accuracy of all the tasks while training them sequentially (T1, T2, and T3) using EWC. It can be noted that at the end of the training, T1 gets an accuracy of 81%, T2 gets 80.3%, and T3 learns to the accuracy of 97%.

**Figure 4.**
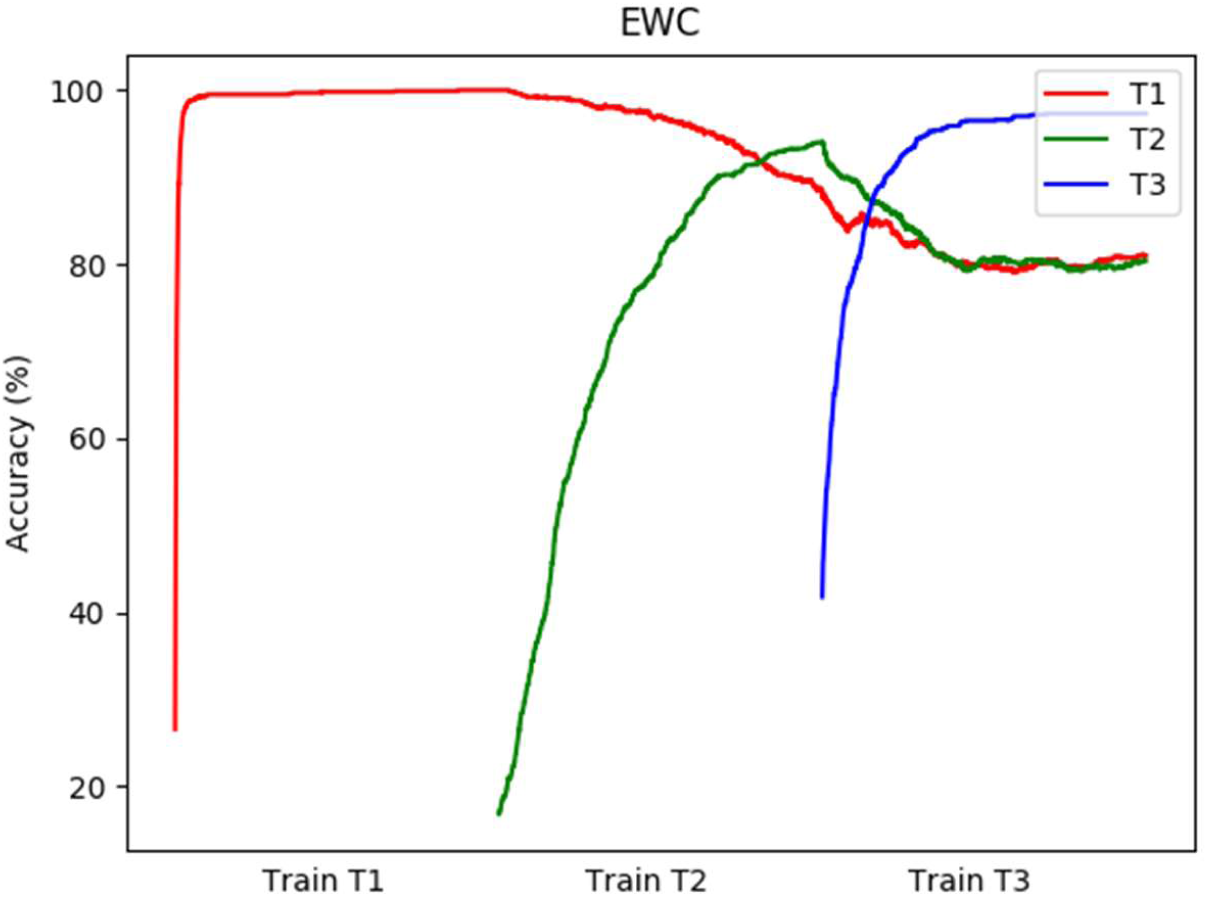
Task accuracies on Split MNIST Dataset using EWC

Figure 5 shows the accuracy of all the tasks while training using the proposed approach on the split MNIST dataset. Here at the end of the training, the tasks T1, T2, and T3 get the accuracy of 88%, 84%, and 92 %, respectively. Here the average accuracy among the three tasks can be observed as 86.1% for EWC and 88% for the proposed model.

**Figure 5.**
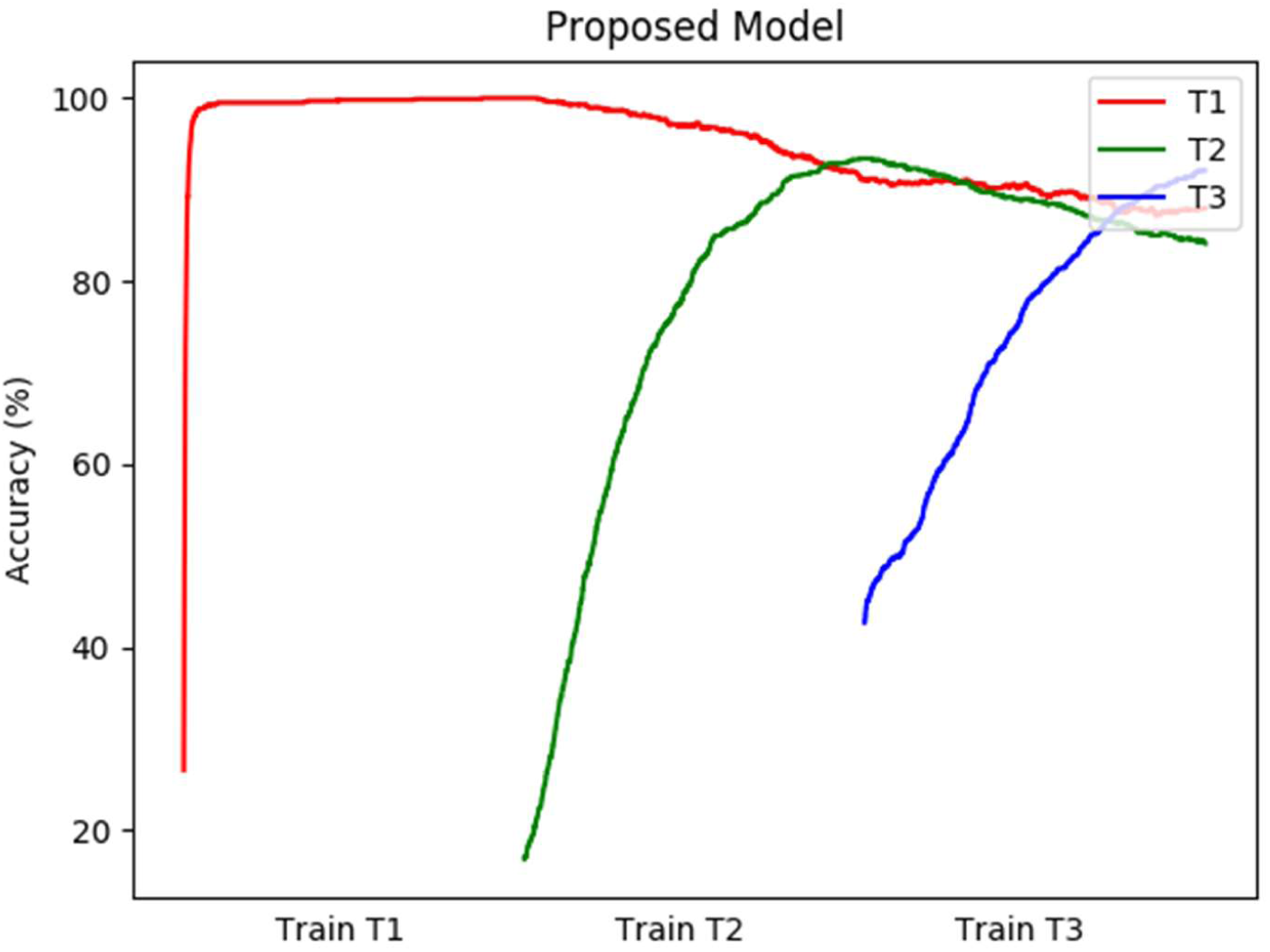
Task accuracies on Split MNIST Dataset using the proposed model

### Convolutional Network

The performance of the above models is evaluated on Convolution-based classification networks using a split MNIST dataset with three tasks (T1, T2, and T3). The network comprises 2 convolution (+max-pooling) layers followed by two fully connected layers. The first convolution layer uses eight 3×3 filters. The second convolution layer uses 16 3×3 filters. The max-pooling layer uses stride two following each convolution layer. The output of the 2^nd^ convolution layer is processed by the fully connected layers with 128 and 3 neurons, respectively. Here both the convolution layers use leaky-ReLU activation function, the first fully connected layer uses sigmoid function, and the output layer uses SoftMax function.

The first task is trained to attain 100% accuracy with L2 regularization, and the performance of the proposed model is compared with EWC while training the second and third tasks sequentially.

Figure 6 shows the cross-entropy loss on tasks T1, T2, and T3 while training with the proposed model. After training with all the tasks, tasks T1, T2, and T3 get the loss .35, .31, and .397, respectively.

**Figure 6.**
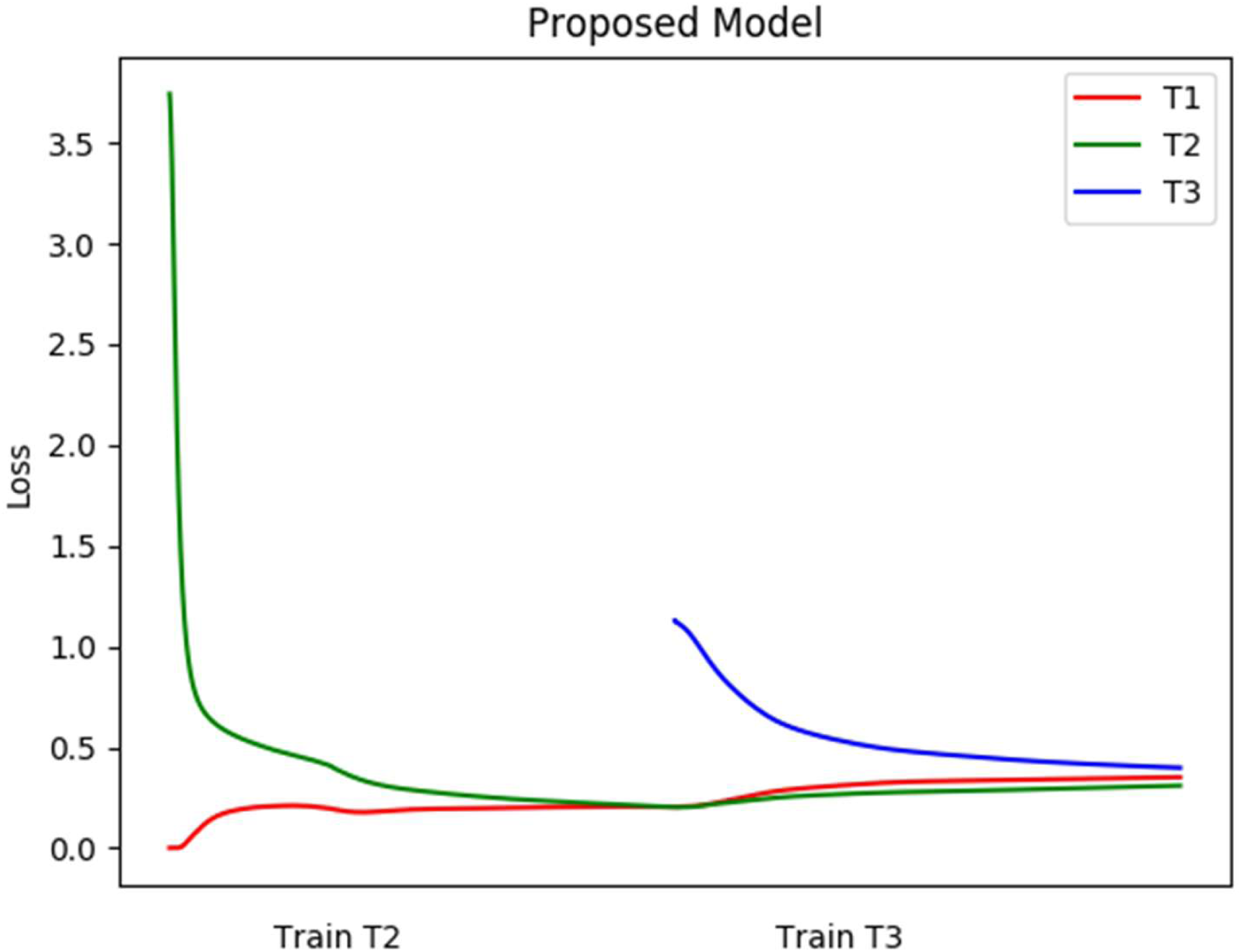
Loss for three Tasks when training with the proposed model on CNN based classifier

Figure 7 shows the loss of the three tasks while training with EWC. After training with all three tasks, the loss for the T1 is retained at .3, T2 at .6, and T3 at .157.

**Figure 7.**
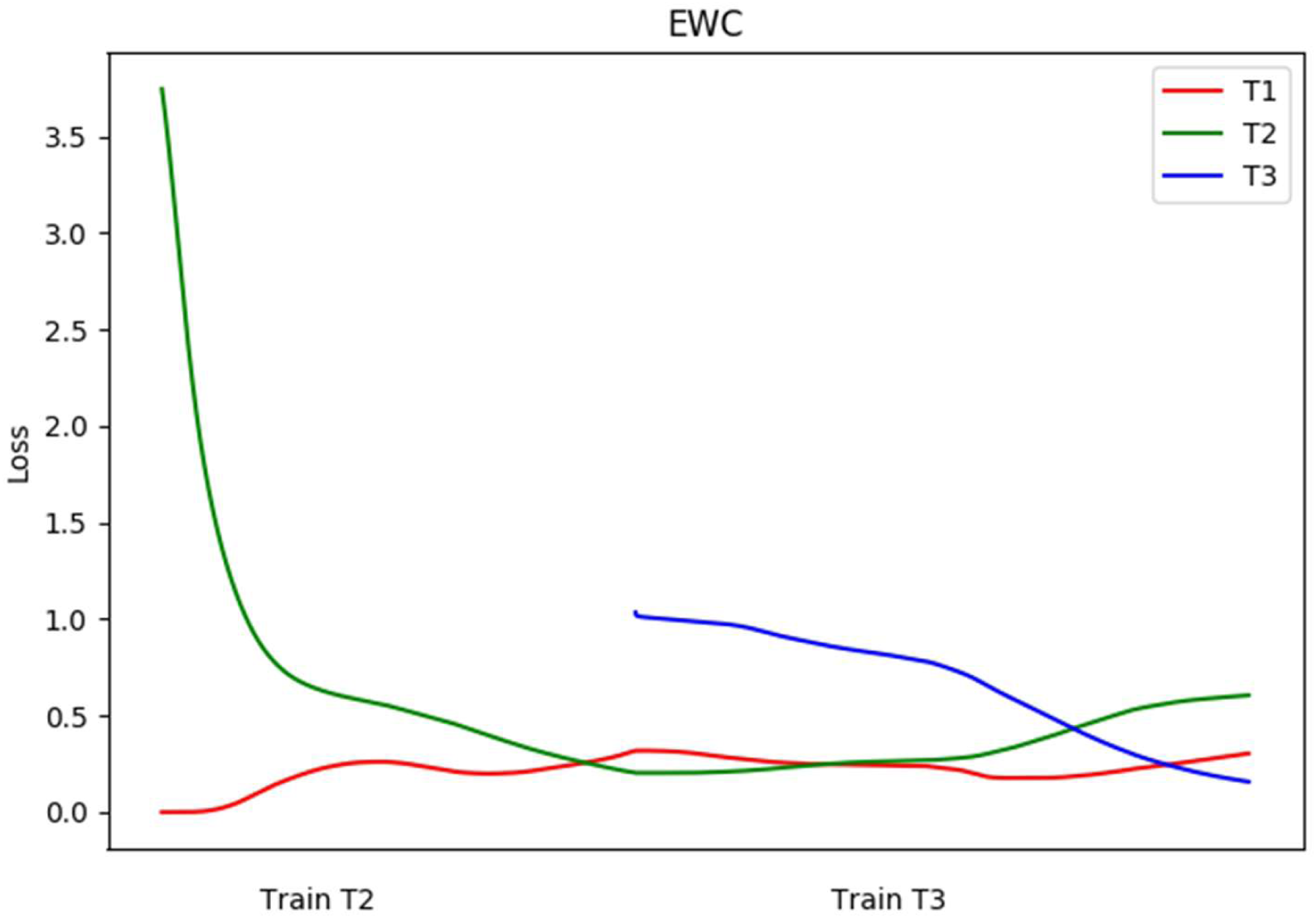
Loss for three Tasks when training with EWC on CNN based classifier

Figure 8 shows the accuracy of the three tasks while training with the proposed model. After training with all the tasks, T1 gets an accuracy of 80.07%, T2 gets 83.67%, and T3 gets 79.3%. For the proposed model, the average accuracy among the three tasks is 81.01%.

**Figure 8.**
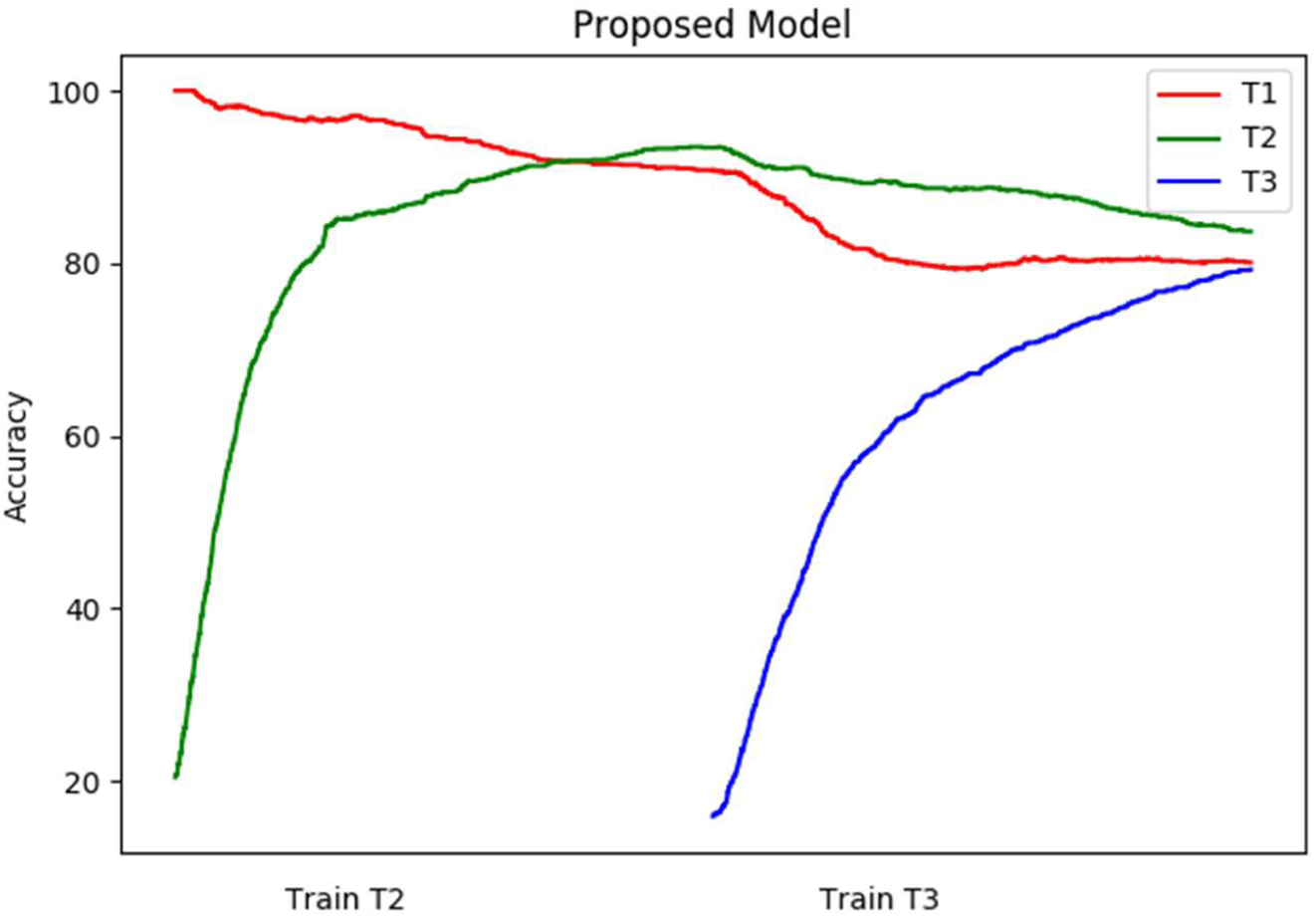
Accuracy for three Tasks when training with the proposed model on CNN based classifier

Figure 9 shows the accuracy of the three tasks when training with EWC. After training, the T1 gets the accuracy of 79%, T2 gets 60%, and T3 gets 90%. The three tasks together get an average accuracy of 76.3% for EWC.

**Figure 9.**
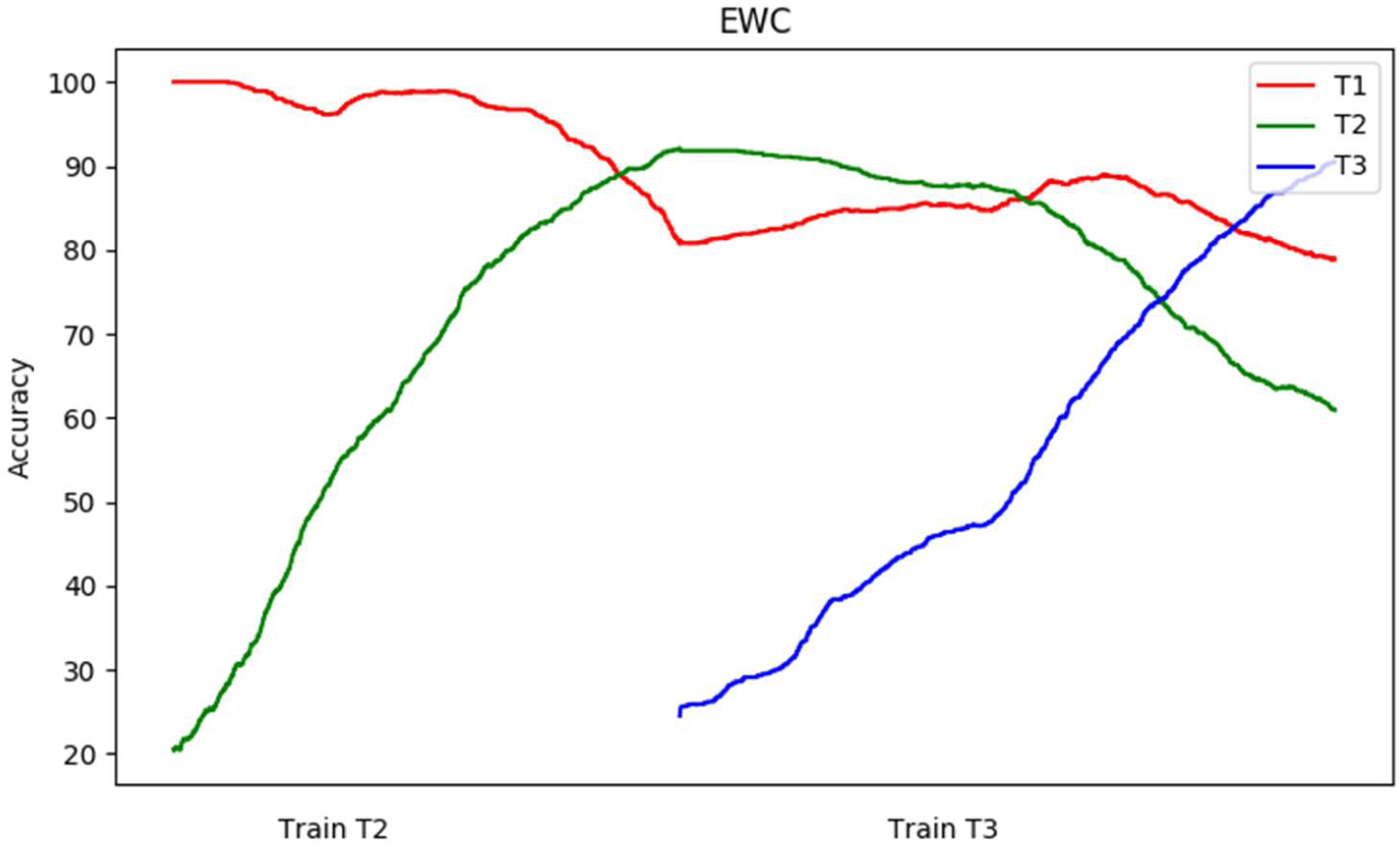
Accuracy for three Tasks when training with EWC

## Auto-Encoder

To evaluate if the proposed model can retain the old information on even more complex networks, we experimented with the continual learning task on an autoencoder. The same split MNIST settings are used here. The number of data points for each task is set to 100.

The autoencoder comprises four fully connected layers. The network receives the MNIST image of size 784. The input is encoded by two fully connected layers with 256 and 128 neurons, respectively. The encoded feature is decoded by two layers with 256 and 784 neurons, respectively. All the layers use the sigmoid activation function. The network is trained to minimize cross-entropy loss. The network is trained using Adam optimizer.

The loss for the three tasks is compared between the proposed model and EWC. Figure 10 shows the cross-entropy loss for each task when training the tasks T1, T2, and T3 sequentially using EWC. At the end of the training, the tasks T1, T2 and T3 get the loss of .126, .122, and .091 with an average task loss of 0.113.

**Figure 10.**
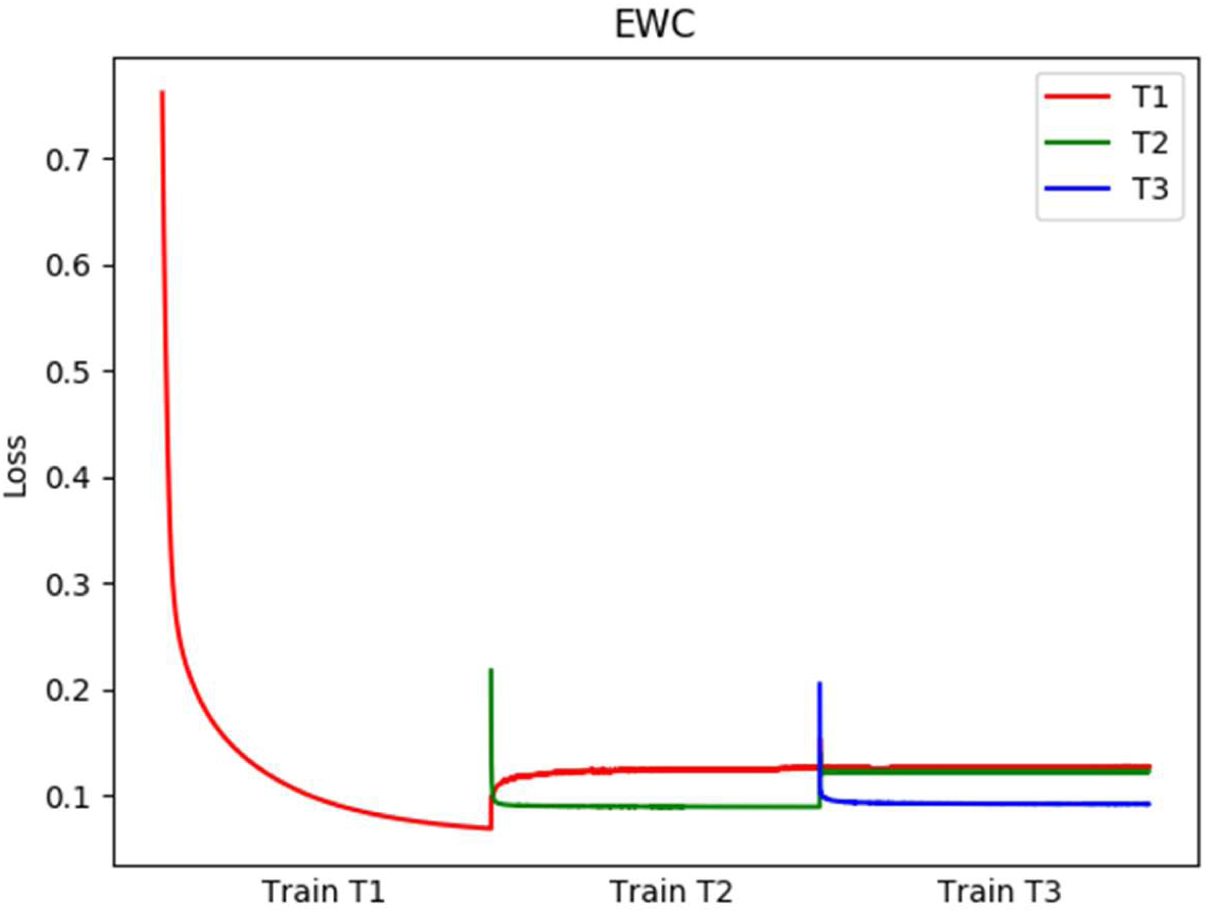
Loss for three Tasks when training with EWC on Autoencoder

Similarly, Figure 11 shows the loss for the proposed method. At the end of the training, T1 gets the loss of .111, T2 gets .123, and T3 gets .078. Here the average loss for the proposed model is 0.104.

**Figure 11.**
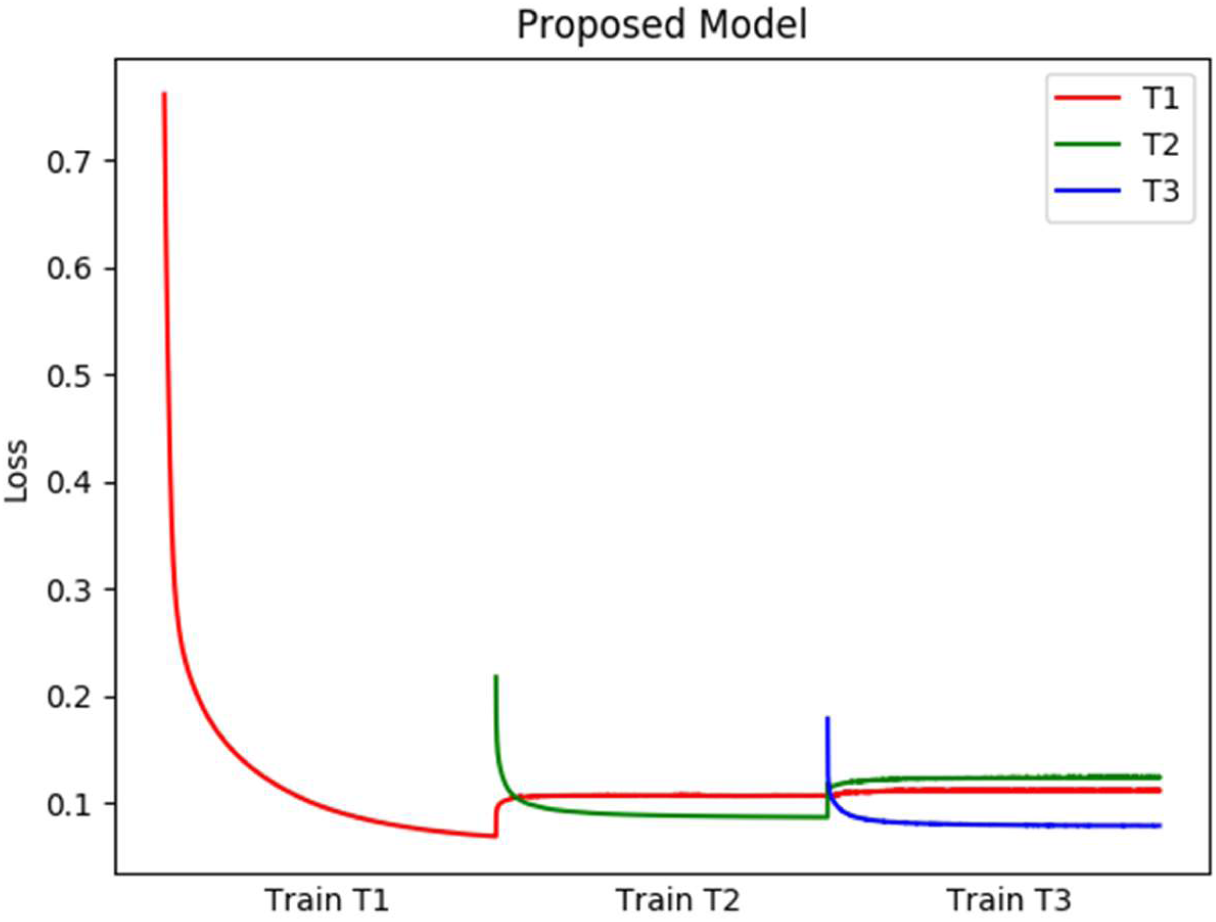
Loss for three Tasks when training with the proposed model on Autoencoder

To visualize the performance of the proposed model on autoencoder, one sample image output from each task at the end of each training stage is shown below.

Figure 12 and Figure 13 displays the sample outputs of the autoencoder while training with EWC and the proposed model, respectively. In both the figures, the row represents the output of the network for one sample image from the particular task. The column represents the end of training each task. At the end of training all three tasks, the proposed model generates a better image with lesser noise patches compared to the EWC.

**Figure 12.**
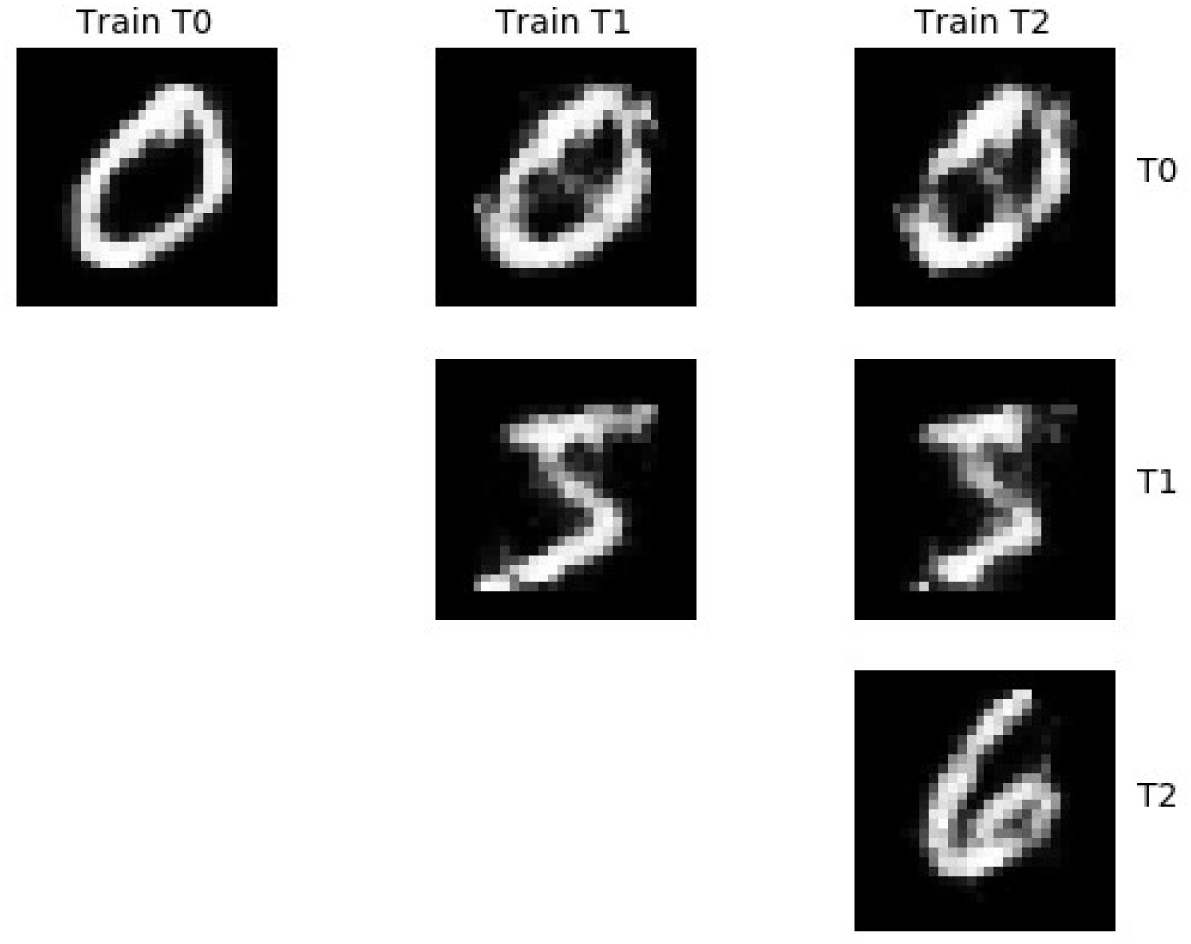
Sample Autoencoder Output from each task while training using EWC

**Figure 13.**
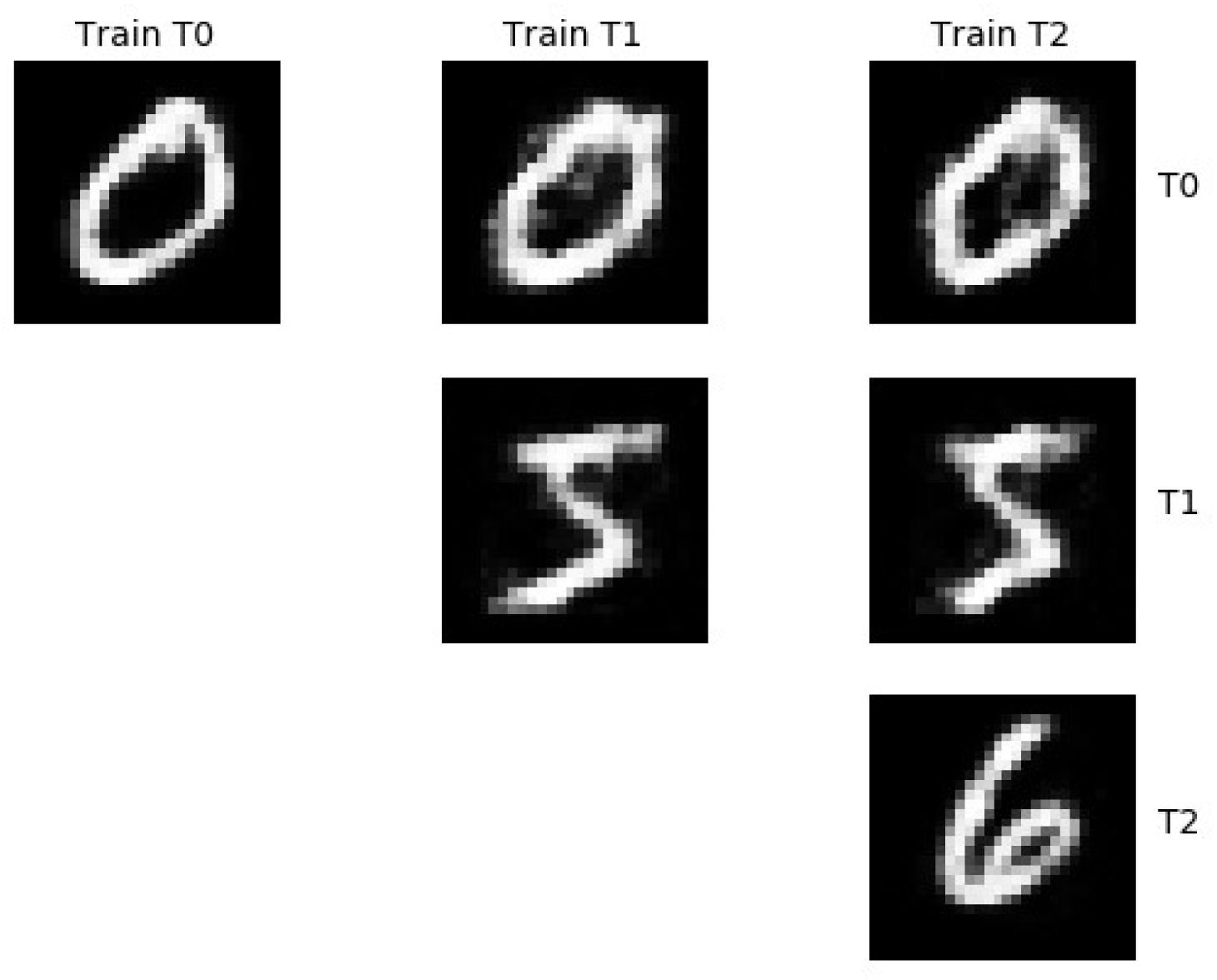
Sample Autoencoder Output from each task while training using the proposed model

### Core50 Dataset

To evaluate the proposed model on complex datasets, CORe50 is used. CORe50 is a complex dataset generated for continual object recognition problems [37]. It contains images of 50 household items under different conditions. The input image is of dimension 128×128×3.

Two types of problems are used for evaluating the performance of the proposed model. In the New instances (NI) case, each task will have the images corresponding to the same classes, but the environment (background, illumination, orientation, etc.) is different between tasks. Here we have considered three tasks, each having 1000 images. Each task consists of images corresponding to 5 classes.

In New classes (NC) scenario, each task will have images corresponding to different types of objects (classes). Each task includes 1000 images corresponding to 5 image classes.

### Network Architecture

The network comprises two convolutional (+max-pooling) layers followed by two fully connected layers. The first convolution layer uses eight 3×3 filters. The second convolution layer uses eight 3×3 filters. The max-pooling layer uses stride 2 following each convolution layer. The output of the 2nd convolution layer is processed by the fully connected layers with 1024 and 5 neurons, respectively. Here both the convolution layers use the leaky-ReLU activation function, the first fully connected layer uses the sigmoid function, and the output layer uses the SoftMax function.

The loss and accuracy of the three tasks are shown while training them sequentially. The first task is trained with standard L2 regularization to get the maximum accuracy of 100%. Figure 14 shows the loss for the three tasks while training them sequentially using EWC. Here at the end of the training, the tasks T1, T2, and T3 get the loss of .988, 1.474, and .218, respectively, with an average loss of .8934.

**Figure 14.**
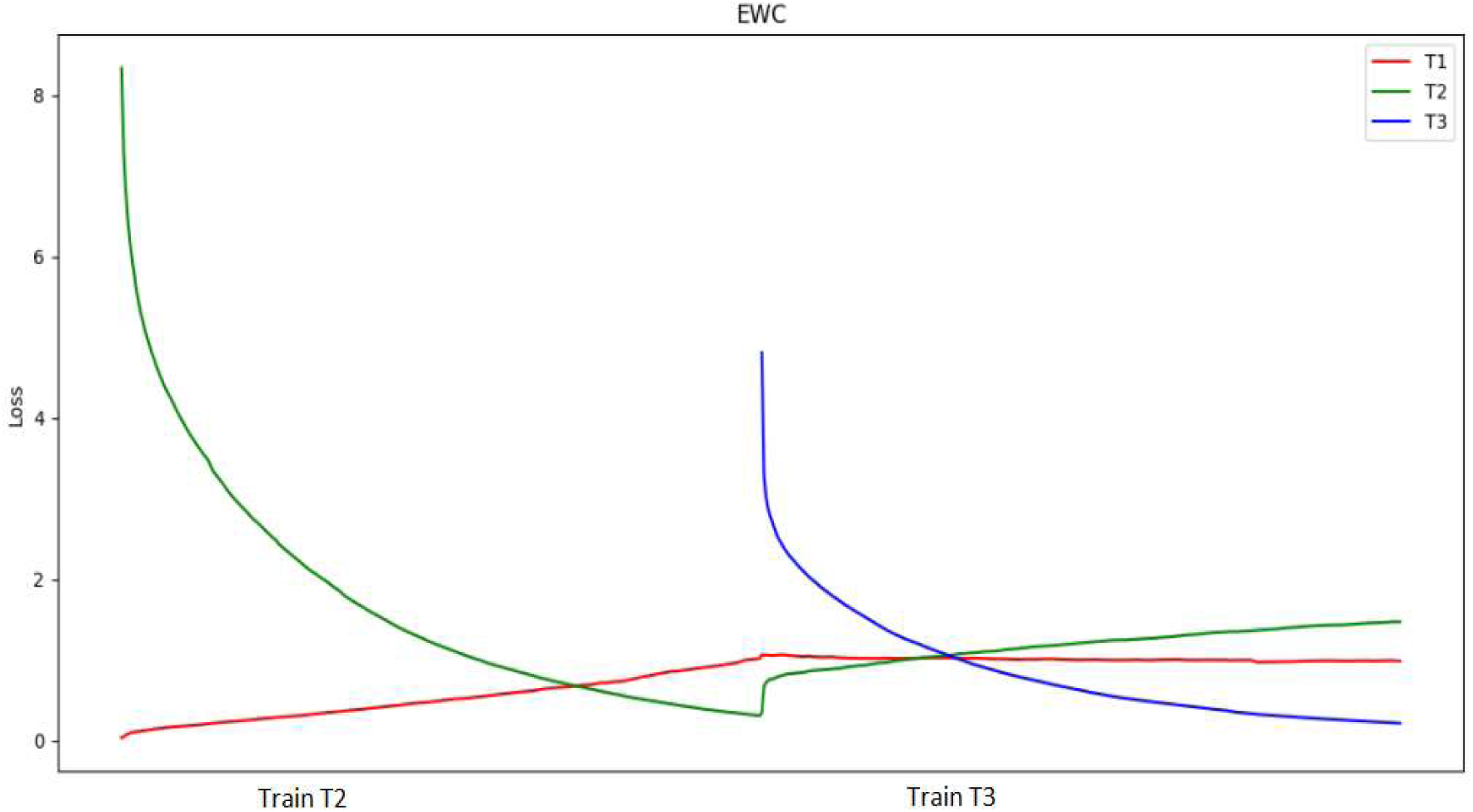
Loss of three New Instances tasks using EWC

Figure 15 shows the accuracy of all three tasks while training with EWC. At the end of training, the three tasks, T1, T2, and T3, get the accuracy of 93.2%, 85.4%, and 100%, respectively, with an average of 92.86%.

**Figure 15.**
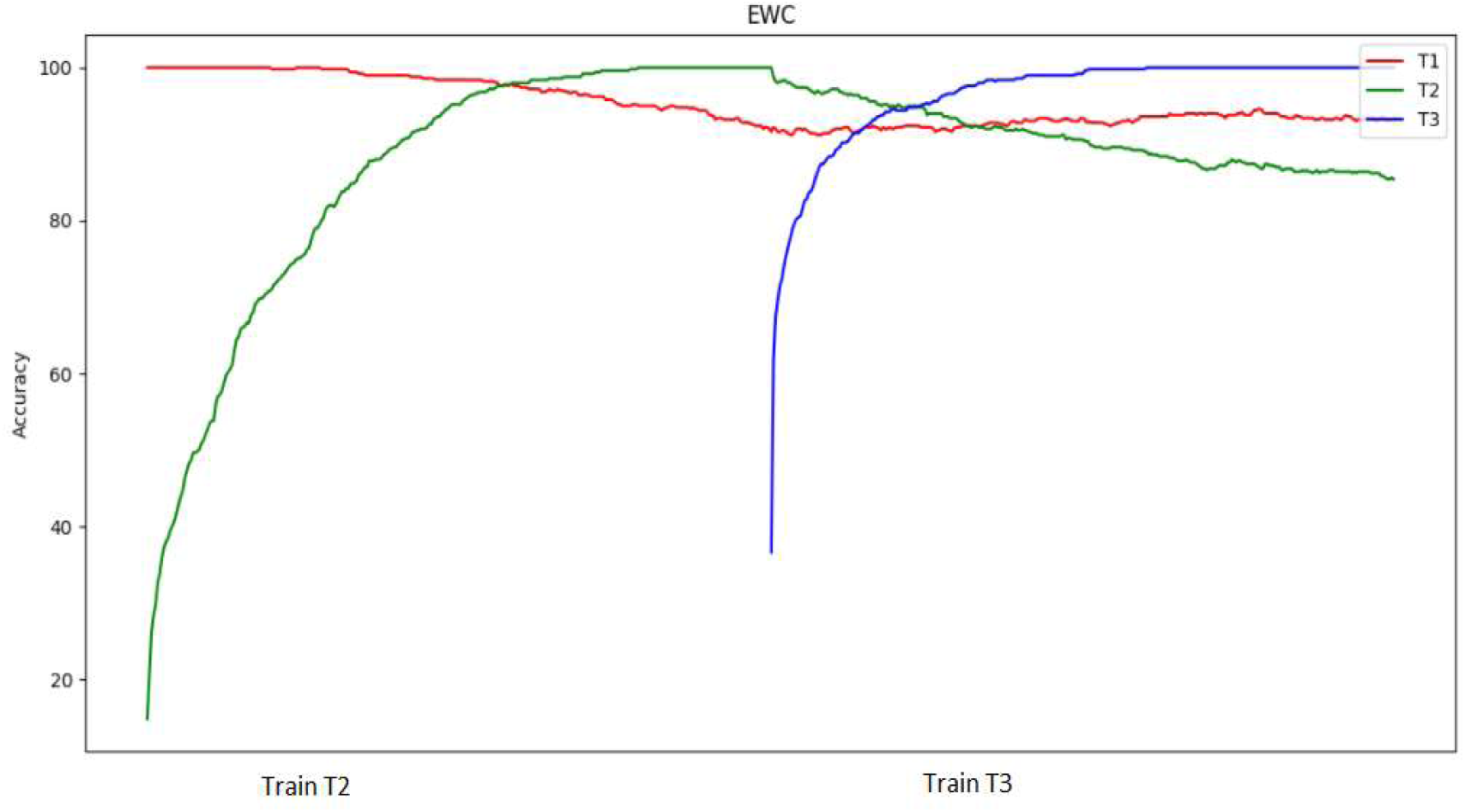
Accuracy of three New Instances tasks using EWC

Figure 16 shows the loss of three tasks while training the network with the proposed model. After training, each task gets the loss of .679, 1.352, and .212 (in order) with an average of .7476.

**Figure 16.**
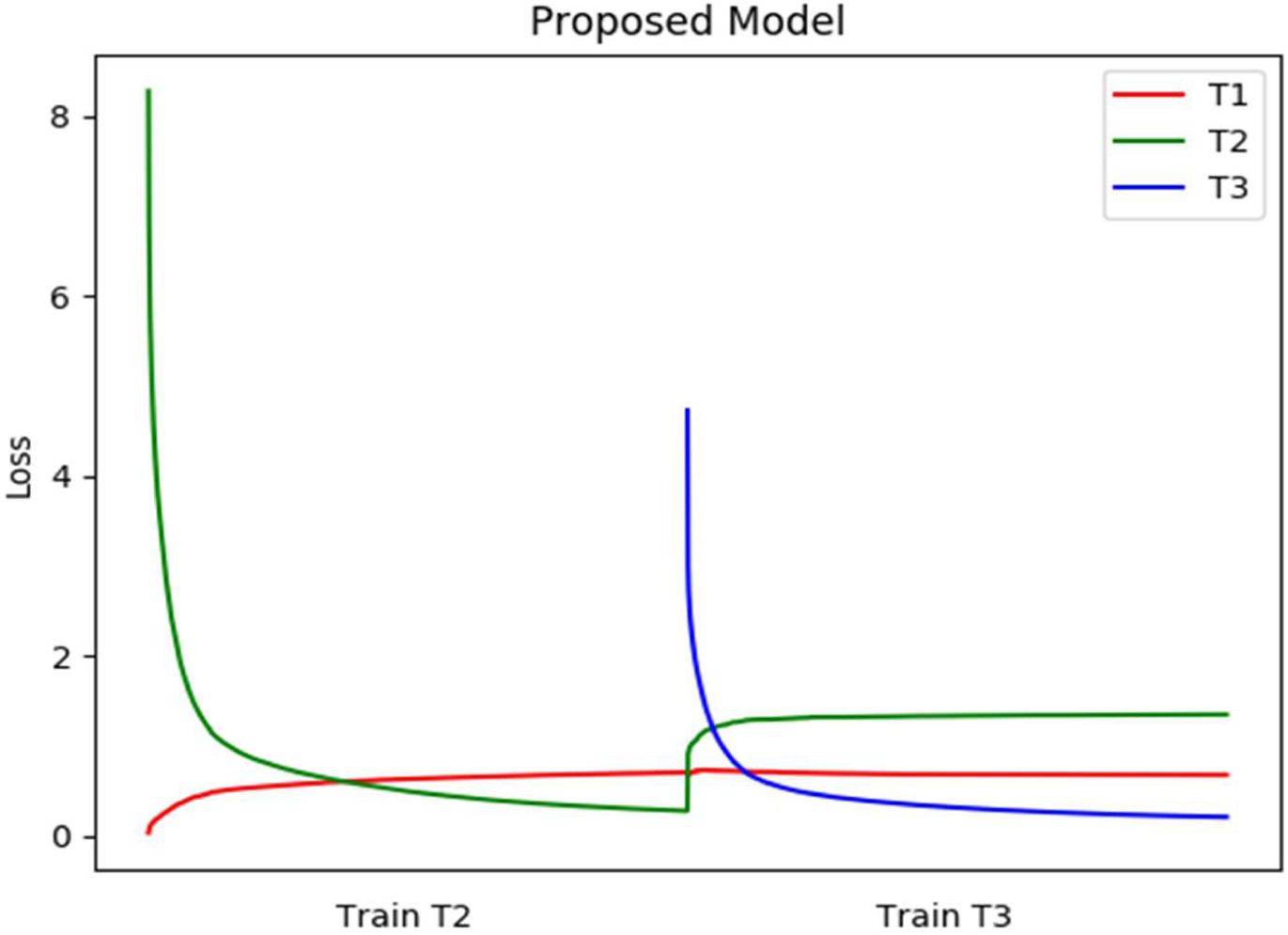
Loss of three New Instance based tasks using the proposed model

Figure 17 displays the accuracy for the three tasks while learning with the proposed model sequentially. Each task gets the accuracy of 96%, 89.2%, and 100% respectively. Here the average accuracy for all the tasks together comes to 95.06%.

**Figure 17.**
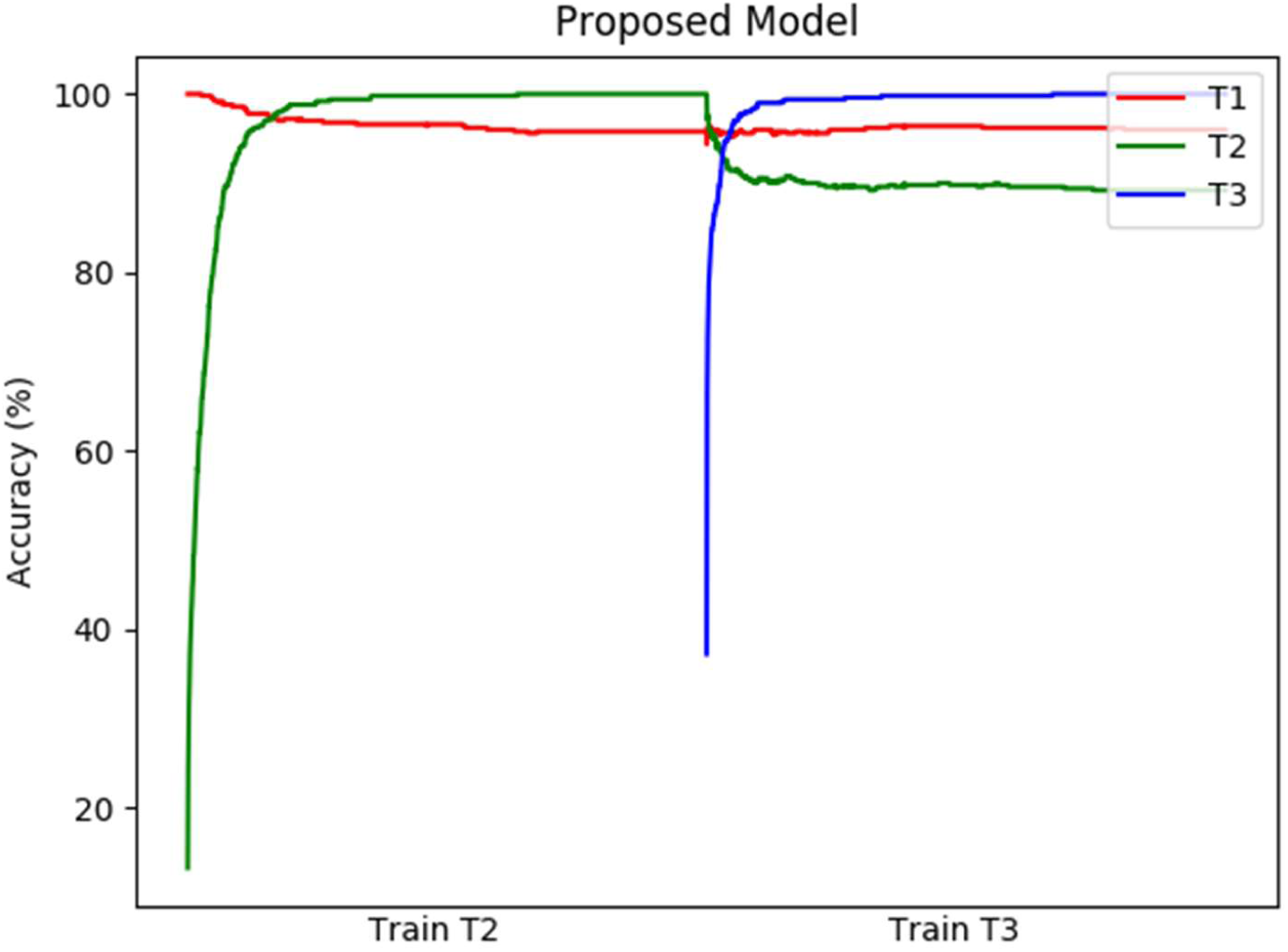
Accuracy of three New Instances tasks using the proposed model

**Figure 18.**
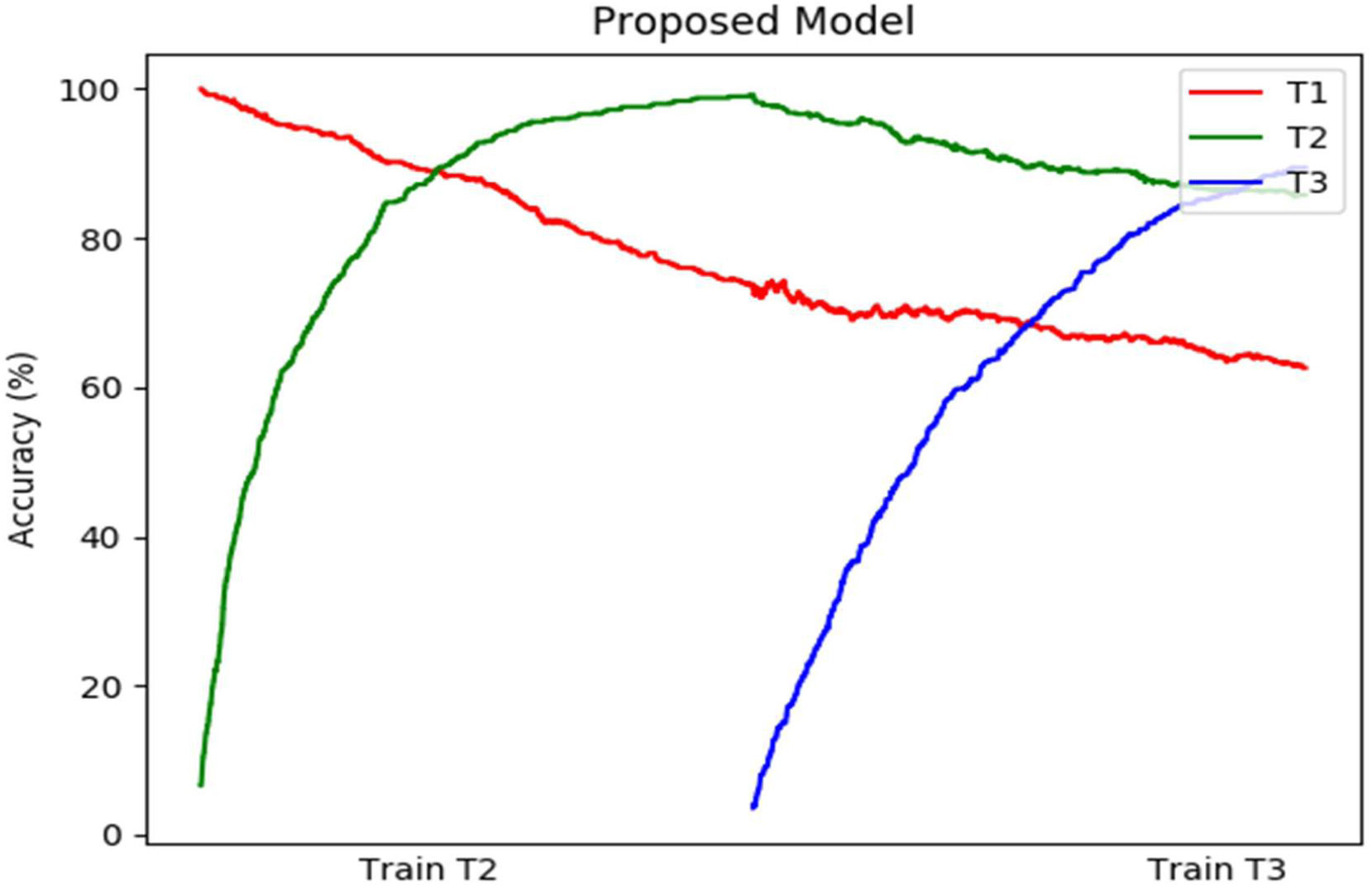
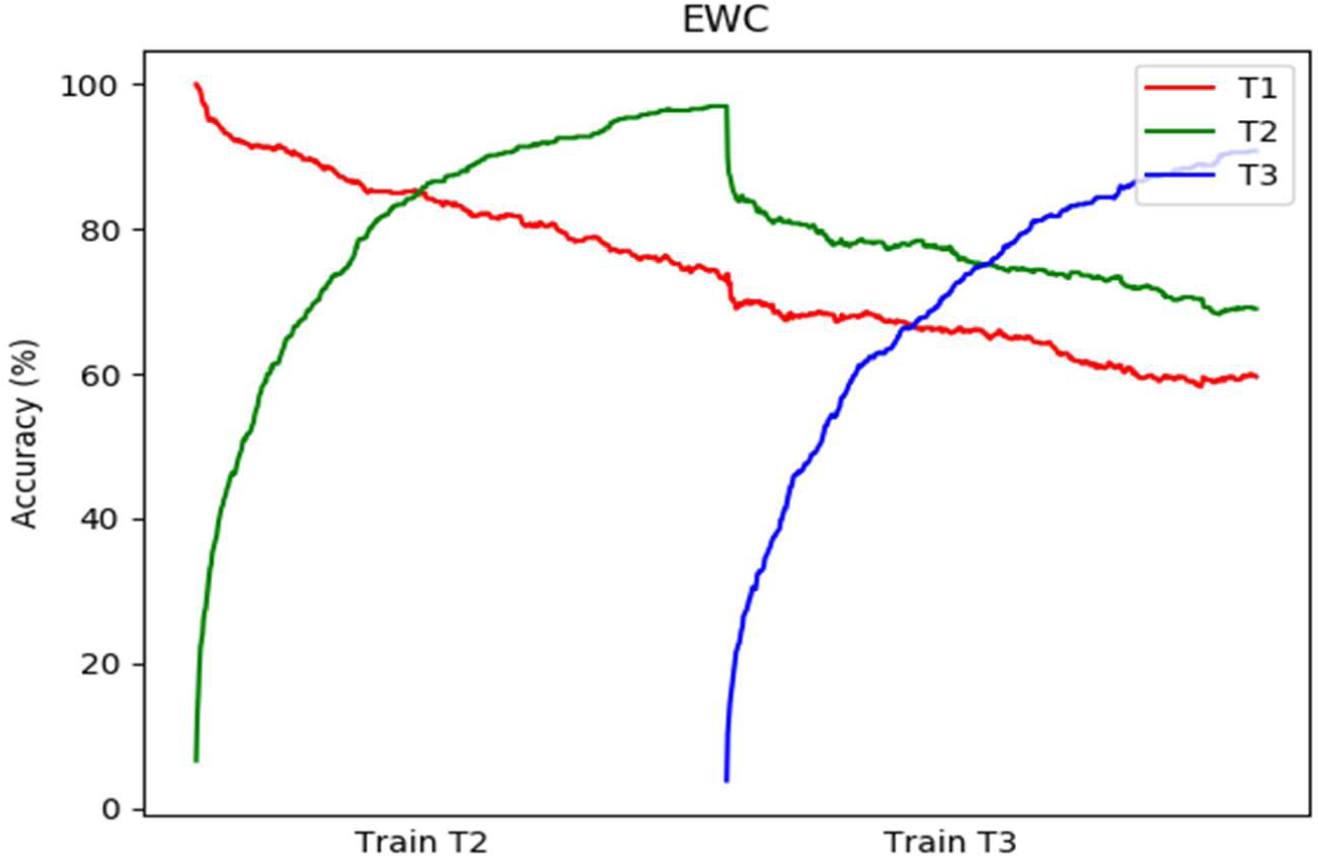
Accuracy of three New Classes task using EWC

Table 2 shows the comparison of average loss and accuracy between EWC and the proposed model on CORe50 new instances task.

**Table 2.** Performance comparison between EWC and the proposed model Core50 New Instances dataset

|  | Average Loss | Average Accuracy (%) |
| --- | --- | --- |
| **EWC** | .8934 | 92.86 |
| **Proposed Model** | .7476 | 95.06 |

Figure 19 shows the accuracy for all the tasks for the New Classes scenario while training with EWC and the proposed model, respectively. In both cases, the first task is trained with the conventional regularization method. The second and the third tasks are trained with the respective approaches. Towards the end of the training, EWC produces the accuracy of 59.6%, 69%, and 90.8% on the three tasks. The proposed model produces the accuracy of 62.6%, 85.8%, 89.4% on the tasks. Here EWC makes an average accuracy of 73.13%, and the proposed model gains the average accuracy of 79.27%.

**Figure 19.**
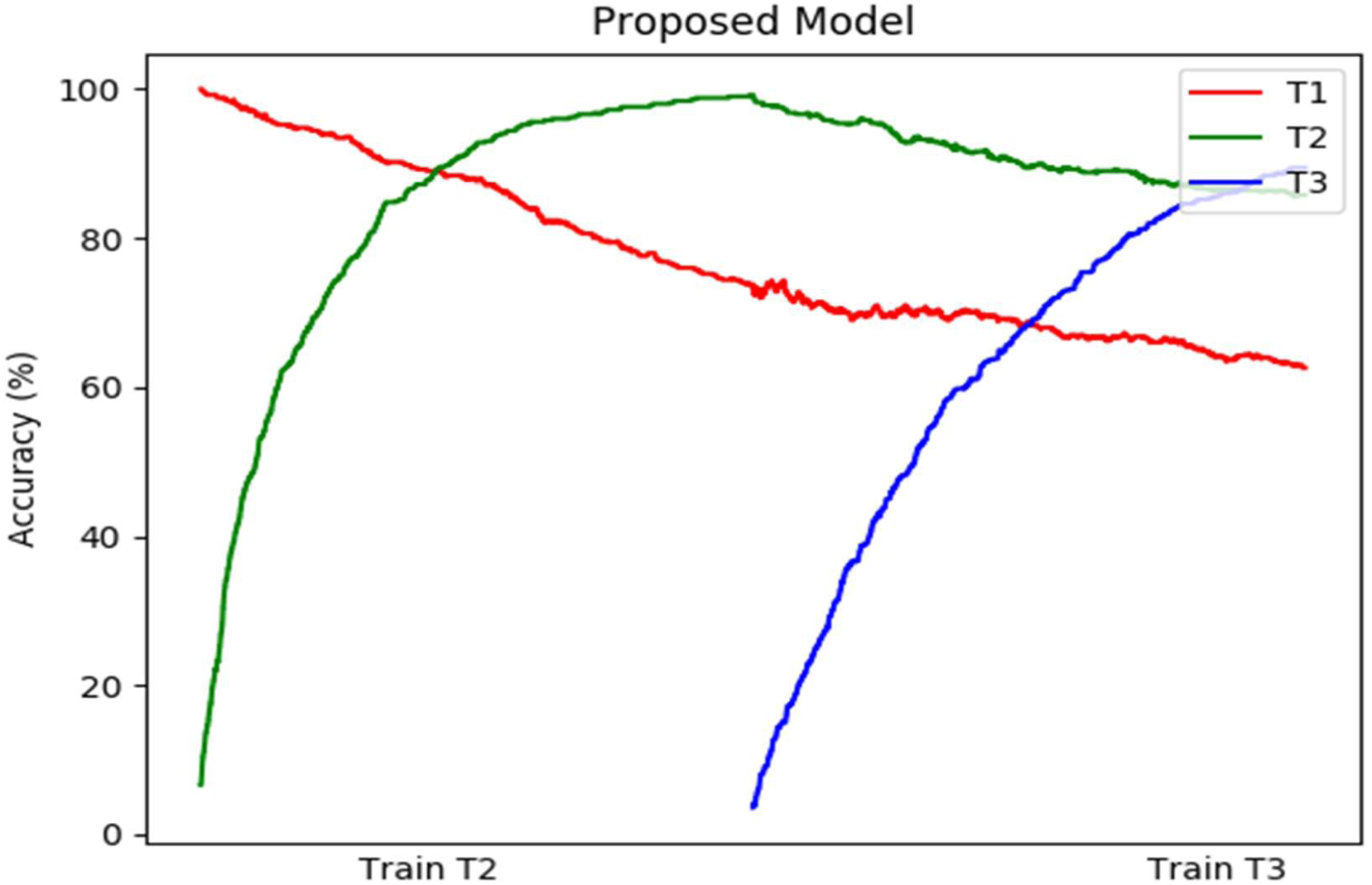
Accuracy of three New Classes task using the proposed model

Similarly, Figure 20 and Figure 21 display the loss for the three tasks while training with EWC and the proposed model, respectively. After training, EWC obtains a loss of 1.897, 1.698, and 1.613 for the three tasks. The proposed model obtains a loss of 1.799, 1.149, and 1.100. EWC makes the average loss of 1.613, and the proposed model gets 1.349.

**Figure 20.**
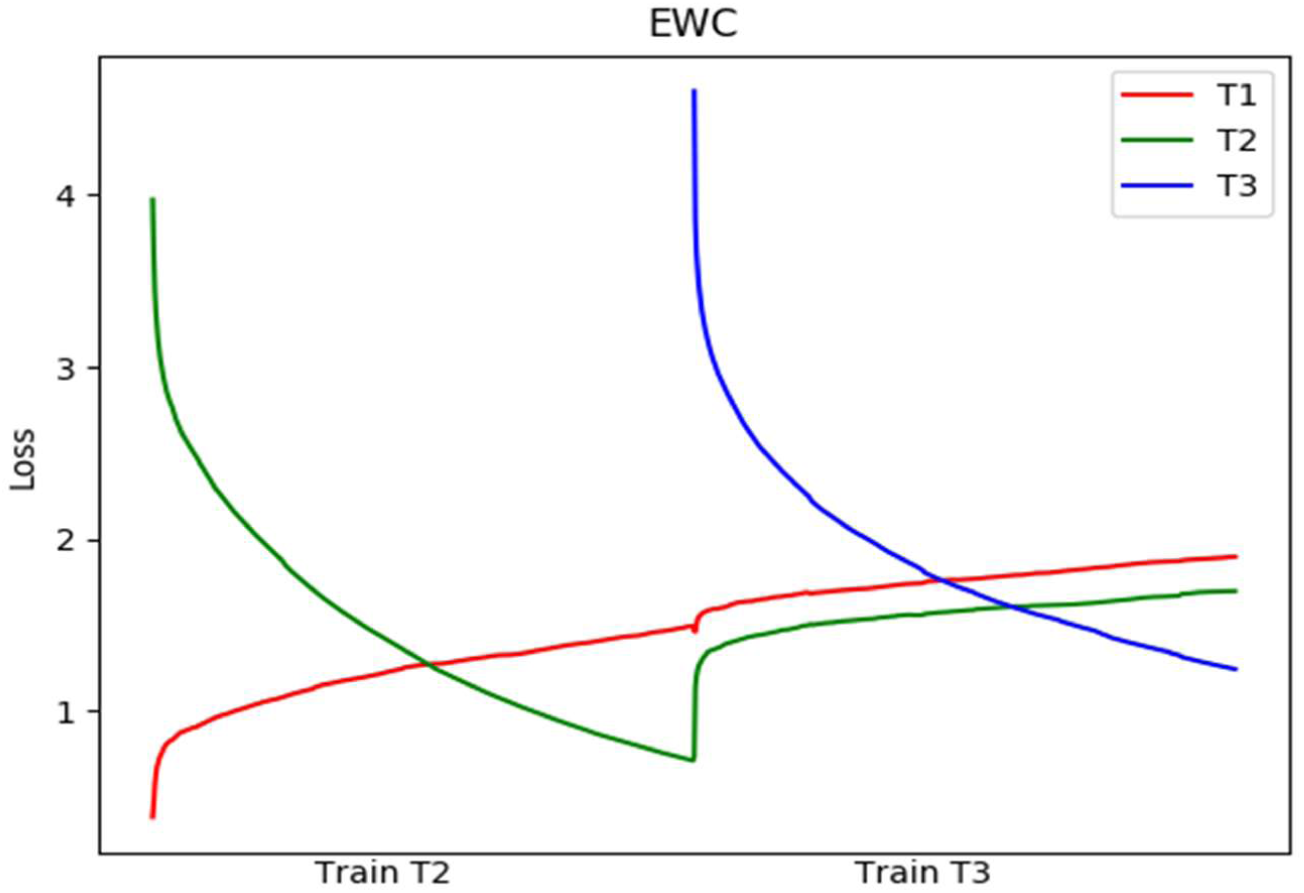
Loss of three New Classes task using EWC

**Figure 21.**
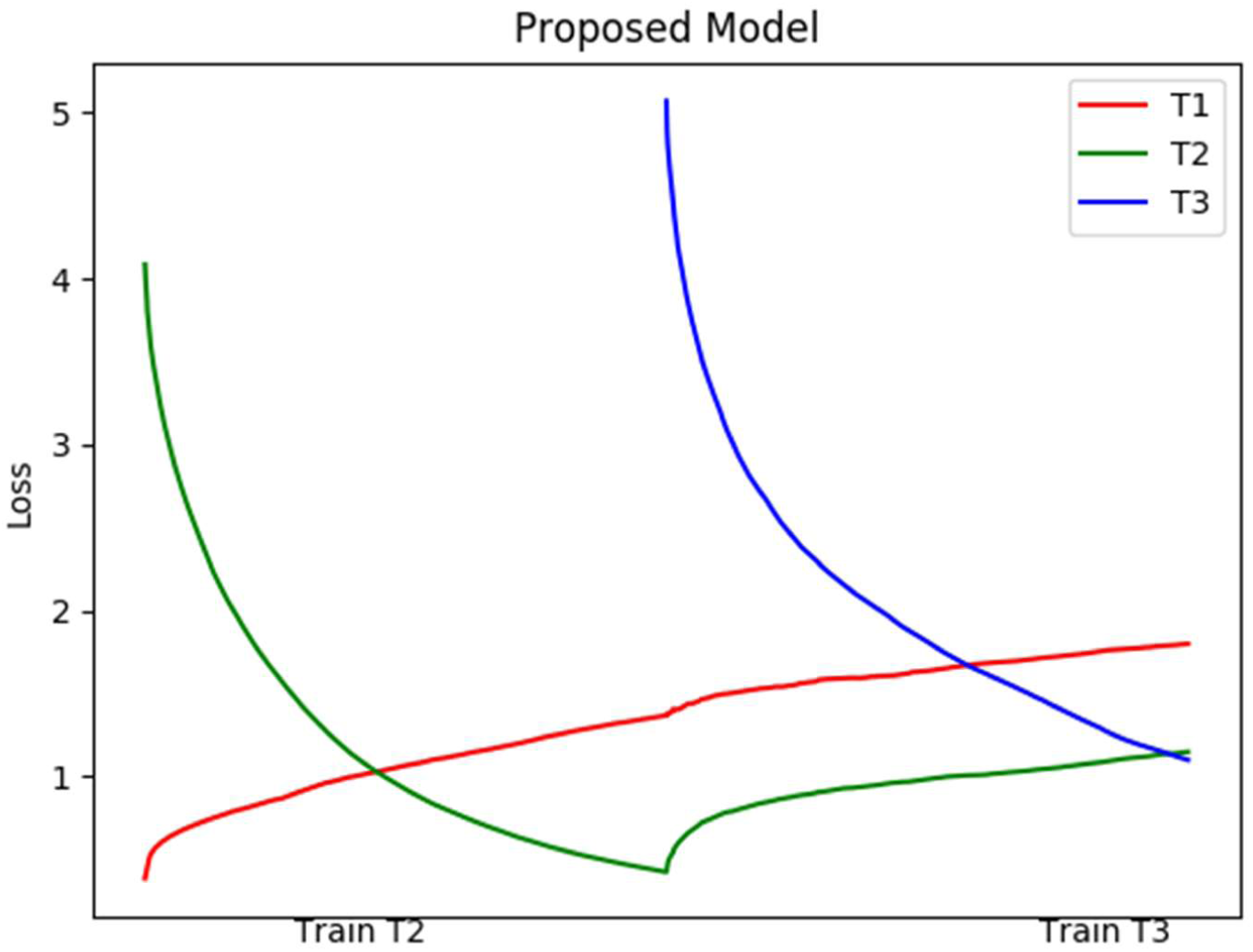
Loss of three New Classes task using the proposed model

Table 3 shows the comparison of average loss and accuracy between EWC and the proposed model on CORe50 new classes task. It may be observed that the proposed model attains a higher accuracy and a smaller loss over EWC.

**Table 3.** Performance comparison between EWC and the proposed model Core50 New Classes dataset

|  | **Average Loss** | **Average Accuracy (%)** |
| --- | --- | --- |
| **EWC** | **1.613** | **73.13** |
| **Proposed Model** | **1.349** | **79.27** |

## Discussion

We present a deep network model of continual learning that acquires and integrates new memories along with the old memories by a process of memory consolidation. The proposed model tries to avoid catastrophic forgetting by imposing a constraint on the network by updating the parameters in a direction orthogonal to the average gradient estimated from the previously learned tasks. This orthogonality constraint is imposed on the parameters at the neuron level.

The performance of the proposed model is compared with the EWC, the state-of-the-art model that uses a similar mechanism. The performance is assessed on various classification tasks such as permuted MNIST (simple task) and split MNIST (simple task), CNN based classification on split MNIST (complex network), and CNN-based classification on CORe50 dataset at two different settings New Instances and New Classes (complex network and complex dataset). The performance is also evaluated on an autoencoder trained on split MNIST dataset (complex network). All the available models on continual learning have been mainly assessed on classification problems. There are no studies in the literature that examine catastrophic interference in autoencoder-based networks. The proposed model is evaluated on autoencoders along with the classification problems. Among the classification tasks, the proposed model shows similar or slightly better performance than EWC on simpler tasks such as permuted MNIST and split MNIST. When the complexity of the dataset or the network increases, the proposed model shows better performance compared to EWC.

## Comparison to Other Models

There are several models that address the catastrophic forgetting problem. The elastic weight consolidation (EWC) approach uses the copy of the diagonals of the Fisher information matrix that is derived from the previously learned tasks. Thus it maintains importance value for each synapse [31]. The higher importance value denotes the importance of a particular parameter for the previous tasks. Similarly, in the synaptic intelligence (SI) approach, the importance value for each parameter is assigned according to the change in the parameter values [32]. In both the above approaches (EWC and SI), the importance is given for each synapse, while in the proposed model, the importance is given for the direction at the neuronal level.

The Self-net approach uses an external autoencoder to represent the parameters for each task in a compressed format [27]. But it did not discuss the parameters used in the autoencoder. The proposed model uses only the copy of the latest trained parameters and the average gradients, which is just two times the number of the network parameters.

The learning-without-forgetting approach uses the current task data to retain the performance on the old tasks [41]. This model uses two sets of parameters: shared parameters and task-specific parameters. However, this kind of task-specific network can be associated with context information. The proposed model uses the same network for different tasks. This association can be introduced by adding contextual information to the network.

In the orthogonal gradient descent method, the predefined number of gradients for each task is maintained, and the updating of parameters for the new task is controlled by updating the parameters orthogonal to all the previously stored gradients [33]. This orthogonal gradient direction is estimated using the Gram-Schmidt method. Here the orthogonality is maintained for the whole set of parameters. Whereas in the proposed model, the orthogonality constraint is restricted to the neuronal level. In the orthogonal gradient descent method, as the number of tasks increases, the size of the copy of the gradients of parameters also increases. In the proposed model, the number of extra parameters does not change with respect to the number of tasks.

Some models use network expansion mechanisms, where at each layer, new neurons are added based on certain criteria [26]. This proposed model can accommodate this kind of network expansion as the orthogonality character can be maintained even when new neurons are added.

The biological plausibility of the proposed model can be briefly explained by comparing it to two biological processes: memory consolidation and synaptic tagging. During initial encoding, the hippocampus stores the recently acquired memory and acts as short-term memory [42]. Memory consolidation process is initiated by the hippocampus replaying the recently acquired memory [13], [14]. This consolidation enables the accumulation of recent memory along with the remote memory in the long-term memory store, and the memory becomes independent from the hippocampus that becomes remote memory. The proposed model does not include the short-term memory and the replay mechanism, which are the characteristics of the hippocampus. These are planned to be included in future studies.

Synaptic tagging is another mechanism that can possibly be related to the proposed model. Frey et al. (1997) proposed that the memory traces set a tag for each synapse that acts as a marker for change in the synaptic plasticity for the consolidation [43]. They also suggest that only the tagged synapses are allowed for consolidation [43]. The synaptic tag denotes the eligibility of a particular synapse for consolidation [44]. Though synaptic tagging was proposed as a mechanism of the neuronal system, recent studies show that astrocytes can play a crucial role in this process. Bosworth et al. (2017) reviewed that the astrocytes are involved in synaptogenesis and synaptic pruning, thereby enhancing the process of complex network formation [45]. Navarette et al. (2012) showed that astrocytes are crucial for cholinergic induced synaptic plasticity and contribute to information storage[46]. Kol et al. (2020) showed the importance of astrocytes in the formation of remote memories [47]. Thus, astrocytes could possibly be the key to this synaptic tagging process. The proposed model uses the copy of the average gradients from the previous tasks and the copy of the latest trained parameters, which can be considered as tags for individual synapses. There is a need for more studies to evaluate this kind of neural mechanism involved in memory consolidation.

## Conclusion

The proposed method gives importance to certain directions while updating the synaptic weights/network parameters (at a neuronal level) rather than giving importance to each synapse (at the synapse level). It helps selectively choose the neurons (whose change in parameters is not orthogonal to the earlier task gradients) and update the parameters corresponding to those neurons faster compared to the other neurons. The proposed model showed better performance in all the tasks while comparing with EWC. This method keeps the copy of the parameters corresponding to the latest trained task and the average gradients for all the previously learned tasks. This provides a possibility of linking the mechanism of synaptic tagging to the parameters used in the neural network models. The model follows the locality principle, where the extra parameters related to a particular neuron alone influence/control the parameter updating. Similarly, the proposed method of memory consolidation provides a theoretical framework to evaluate memory formation and retention in the form of long-term memory. For future work, one possible direction is that instead of using the gradients’ average, the gradients’ important direction can be identified using PCA at the neuron level, and it could improve the performance. Another possible solution is imposing sparsity constraints on the parameters and evaluating the performance.

## Author Contributions

Simulations were done by TK and RK. The main text is written by TK. VSC contributed in providing the key ideas, editing the manuscript drafts, providing insight into the model. BR contributed in providing key ideas, correcting the manuscript drafts. RM contributed in providing key ideas, correcting the manuscript drafts.

